# Differential role of phosphorylation in glucagon family receptor signaling revealed by mass spectrometry

**DOI:** 10.1101/2025.03.10.642457

**Authors:** Ian M. Lamb, Alex D. White, Francis S. Willard, Michael J. Chalmers, Junpeng Xiao

## Abstract

In response to extracellular ligands, G protein-coupled receptors (GPCRs) undergo conformational changes that induce coupling to intracellular effectors such as heterotrimeric G proteins that trigger various downstream signaling pathways ^1^. These events have been shown to be highly regulated by the concerted effects of post-translational modifications (PTMs) that occur in a ligand-dependent manner. Most notably, phosphorylation of residues in the C-terminal cytoplasmic tail of GPCRs has been strongly implicated in promoting receptor interactions with β-arrestins (βarrs) ^2, 3^ which are cytosolic adaptor proteins that modulate G protein coupling ^4^, receptor internalization ^5^, and perhaps also serve as signaling modules in their own right ^6^. Here, we use proteomic methods to identify C-tail residues that are phosphorylated in the glucagon family of class B1 GPCRs (GLP-1R, GCGR, GIPR) upon agonist addition. Mutagenesis studies reveal unique effects of phosphorylated residues on βarr recruitment and cyclic AMP (cAMP) production that occur in a receptor-dependent manner. We demonstrate that phosphorylation of GLP-1R and GIPR is a critical determinant in driving the formation of GPCR-βarr complexes. However, our results suggest that ligand-induced βarr recruitment to the GCGR proceeds in a phosphorylation-independent manner. These findings highlight the importance of recognizing phosphorylation as a component in the regulation of class B1 GPCR signaling, but also the need to consider how such phenomena may not necessarily yield identical effects on intracellular signaling cascades.

## Introduction

The glucagon receptor family of G protein-coupled receptors (GPCRs) is comprised of six members that serve critical roles in proper energy and nutrient homeostasis. Of these, the cognate receptors for glucagon (GCG), glucagon-like peptide-1 (GLP-1), and gastric inhibitory peptide (GIP), or GCGR, GLP-1R, GIPR, respectively, have garnered widespread attention in the last decade as promising therapeutic targets by the pharmaceutical industry for both the treatment of type two diabetes as well as obesity ^7^. Indeed, numerous mono-agonists targeting the GLP-1R including semaglutide ^8^, dulaglutide ^9^, and exenatide ^10^ are FDA approved drugs that have shown remarkable success in the treatment of type 2 diabetes mellitus. Additionally, the dual agonist tirzepatide (targeting GLP-1R and GIPR) was recently approved for the treatment of obesity ^11^ and type 2 diabetes ^8^. Furthermore, the triple agonist retatrutide (targeting GLP-1R, GIPR, and GCGR) has shown remarkable effects on weight loss and is currently in advanced clinical trials ^12^.

As members of the class B1 subfamily of GPCRs, the GCGR, GLP-1R, and GIPR bind and mediate the physiological actions of endogenous peptide ligands. In a cellular context, activation of these receptors is foremost associated with coupling to heterotrimeric G_s_ proteins and production of cyclic adenosine monophosphate (cAMP) via activation of adenylyl cyclase ^1^. In addition to G_s_ signaling, these receptors are also associated with the recruitment of β-arrestins (βarrs), which are cytosolic adaptor proteins that are classically considered to both sterically block further G protein coupling as well as mediate translocation of the ligand-receptor complex to endocytic compartments to block subsequent rounds of receptor activation by extracellular ligand ^13^. Mechanistically, the recruitment of βarrs is thought to be preceded by ligand-induced phosphorylation of the intracellular C-terminal tail (hereafter, C-tail) residues by GPCR kinases (GRKs), constituting the so-called “phosphorylation barcode” hypothesis in GPCR signaling ^3, 14^. These phosphorylation motifs have thus been proposed to be the main determinants in driving interactions between activated receptor and βarrs. While a role for cAMP production by each of these receptors has been shown as a primary determinant in driving the physiological and therapeutic actions of these receptors, whether these responses are shaped upon interaction with βarrs remains poorly understood. Furthermore, the requirement of glucagon receptor family phosphorylation in driving βarr recruitment and subsequent downstream signaling remains largely unexplored.

Traditional methods for detecting GPCR phosphorylation, such as radioactive orthophosphate labeling and use of phosphorylation-specific antibodies generally lack the ability to identify site-specific protein post translational modifications (PTMs) as well as quantify the magnitude of change that occurs upon receptor activation ^15, 16^. In contrast, advances in mass spectrometry (MS) analytical approaches have enabled the precise localization and quantification of PTMs including phosphorylation of GPCRs ^17–20^ which has historically proven difficult for membrane proteins ^21^, especially GPCRs ^17–19^. However, there are rather few examples of in-depth analyses regarding GPCR phosphorylation and its consequences on signaling cascades and biology. To this end, we have harnessed the capabilities of MS-based PTM detection in combination with optical and biochemical techniques to provide insight into the nature of ligand-induced phosphorylation and its consequences on the signaling properties of GCGR, GLP-1R, and GIPR.

### Experimental Procedures

#### Materials

All peptides were obtained or synthesized at >95% purity. GIP(1-42) (4030658) and GLP-1(7-36) (4030663) peptides were obtained from Bachem. GCG(1-29) was synthesized at Eli Lilly. A biotinylated GCGR agonist was synthesized with the amino acid sequence His-Aib-Gln-Gly-Thr-Phe-Ile-Ser-Asp-Lys(Biotin_PEG4)-Ser-Lys-Tyr-Leu-Asp-Aib-Arg-Ala-Ala-Gln-Asp-Phe-Val-Gln-Trp-Leu-Met-Asp-Thr (CPC Scientific). A biotinylated GCGR antagonist was synthesized with the amino acid sequence (S)_2_Hydroxy_3_phenylpropanoyl-Thr-Ser-Asp-Lys(Biotin_PEG4)-Ser-Lys-Tyr-Leu-Asp-Ser-Arg-Arg-Ala-Gln-Asp-Phe-Val-Gln-Trp-Leu-Met-Asp-Thr-NH2 and a non-biotinylated GCGR agonist was synthesized with the amino acid sequence (S)_2_Hydroxy_3_phenylpropanoyl-Thr-Ser-Asp-Lys(Palm_gammaGlu_gammaGlu)-Ser-Lys-Tyr-Leu-Asp-Ser-Arg-Arg-Ala-Gln-Asp-Phe-Val-Gln-Trp-Leu-Met-Asp-Thr-NH2 (CPC Scientific) ^37^.

#### Cell culture and Transient Transfections

FreeStyle^TM^ 293-F cells (ThermoFisher Scientific, #R79007) were grown in suspension using FreeStyle^TM^ 293 Expression Medium (ThermoFisher Scientific, #12338018). Transfection for pharmacology experiments were conducted as previously described using Fugene-6 (Promega) ^38^. For proteomics experiments, 4.8mL of FreeStyle^TM^ 293 Expression Medium (ThermoFisher Scientific, #12338018) was pre-warmed to 37°C and added to a 15mL conical tube. Subsequently, 240 µL of room-temperature FuGENE ® 6 (Promega, #E2691) were added to the medium, mixed by inversion, and allowed to rest at room temperature for five minutes. 40 µg of the plasmid of interest was added to the FuGENE ® 6 containing medium, mixed by inversion, and allowed to rest at room temperature for 15-20 minutes. The plasmid/ FuGENE ® 6/medium mixture was pipetted into 160mL of FreeStyle^TM^ 293-F cells at a density of 0.25E^6^ cells/mL in a 250 mL shaker flask and placed in a shaking incubator for 48 hours at 37°C.

#### Sample generation for proteomics assays

FreeStyle^TM^ 293-F cells in a 500mL shaker flask in 160mL of FreeStyle^TM^ 293 Expression Medium (ThermoFisher Scientific, #12338018) at a density of 0.25E^6^ cells/mL were transfected with the GPCR construct of interest (GPCR-FLAG for bottom-up proteomics workflows or GPCR-TEV-FLAG for middle-down proteomics workflows). Forty-eight hours later, cells were pelleted by 5-minute centrifugation at 1,000 rpm and washed once with 20mL PBS (Gibco, #14-190-144). Cells were resuspended in 40 mL FreeStyle^TM^ 293 Expression Medium and 20 mL of the suspension was placed into 2 x 125 mL shaker flasks. For stimulation with endogenous peptide ligands, GLP-1 (7-36), GIP (1-42), or GCG (1-29), were solubilized in DMSO (Sigma Aldrich, #D8418) and added to the flask to a final concentration of 1 µM. Synthetic GCG-derived biotinylated peptide ligands were also added to a final concentration of 1µM (supplemental Figure 1C). The other flasks received an equal volume of DMSO as a control and both flasks were placed into a shaking incubator at 37°C for 10 minutes. Cell suspensions from each flask were spun down at 1,000 rpm in 50mL conical tubes, washed once with 20mL of ice-cold PBS, and placed on ice. For all bottom-up proteomics experiments, two milliliters of lysis buffer (1% Triton X-100, 25mM Tris, pH7.5/150mM NaCl, 1mM EDTA/1mM EGTA, UltraPure distilled water) containing 1X HALT protease and phosphatase inhibitor (ThermoFisher Scientific, #1861281) were added to each tube and mixed by pipetting following ligand treatment. Tubes were allowed to rest on ice for 15 minutes to lyse. Lysate was collected by spinning at 14K rpm for 20 minutes at 4°C. For experiments using endogenous peptide ligands, samples were quantified by BCA assay (ThermoFisher Scientfic, #23225 and #23227). For experiments using synthetic biotinylated peptide ligands, no quantification was performed and 250 µl of lysate from each sample was loaded directly onto streptavidin-conjugated AssayMap Bravo tips (see immuno-precipitation section). For middle-down proteomics experiments, after stimulating and washing, pellets were re-solubilized in 5mL of ice-cold 50mM Tris-HCL pH 7.5 (Invitrogen, #15567-027) containing cOmplete^TM^ protease inhibitor cocktail (Roche, #11836145001) and allowed to rest on ice prior to being subjected to a membrane preparation protocol and subsequent quantification by BCA assay.

#### Membrane preparation (middle-down proteomics workflow)

For middle-down proteomics experiments, after ligand treatment and washing, pellets were resuspended in 5mL of ice-cold 50mM Tris-HCL pH 7.5 (Invitrogen, #15567-027) containing cOmplete protease inhibitor cocktail (Roche, #11836145001) and allowed to rest on ice. Cell suspension was homogenized with a glass homogenizer (Wheaton USA) using 15-20 strokes with an overhead motorized drive (PALMGREN 10” drill press, #80110A). Homogenate from each sample was poured into a 50mL conical tube and spun at 2,700 rpm in a Beckman Allegra X-14R centrifuge for 10 minutes at 4°C. Supernatant from each sample was collected and allowed to rest on ice. The pellet for each sample was then resuspended in 5mL of ice-cold 50mM Tris-HCL pH 7.5 containing cOmplete protease inhibitor cocktail and the homogenization step was repeated. After repeating the centrifugation step, supernatant was again collected and combined with the initial supernatant from each sample (two full rounds of homogenization). The remaining pellet was discarded. Combined supernatant from each sample was then poured into Beckman round bottom centrifuge tubes (Ref #357005) and spun 35k x g for 1 hour at 4°C in an Avanti JXN-26 centrifuge (Beckman Coulter) using a JA-20 fixed-angle rotor (Beckman Coulter). Supernatant was decanted and remaining pellet was re-suspended in 250µL lysis buffer (1% Triton X-100, 25mM Tris, pH7.5/150mM NaCl, 1mM EDTA/1mM EGTA, Ultra-Pure distilled water) containing 1X HALT protease and phosphatase inhibitor (ThermoFisher Scientific, #1861281). Pellets were mixed via pipetting, transferred to 1.5mL tubes, and rotated at 4°C for 15 minutes. Each sample was spun for 10 minutes at max speed at 4°C using an Eppendorf 5430R centrifuge, and the supernatant was collected for BCA quantification.

#### TEV protease cleavage reactions

For middle-down proteomics experiments, after membrane preparation and BCA quantification, 100µg of lysate per sample (4 samples per treatment group; 8 samples for one individual experiment) was subjected to TEV cleavage. The appropriate volume of each sample in lysis buffer (1% Triton X-100, 25mM Tris, pH7.5/150mM NaCl, 1mM EDTA/1mM EGTA, UltraPure distilled water) was pipetted into a 1.5mL micro-centrifuge tube along with 25.5µl of 0.1M DTT (10mM final concentration, provided with AcTEV^TM^ protease kit) and 80U of AcTEV^TM^ protease (ThermoFisher Scientific, #12575015). Lysis buffer was added to the tube to bring up the final volume to 255µL. Tubes were rocked gently overnight at 4°C and the entire reaction was subjected to immuno-precipitation the following day.

#### Receptor Immuno-precipitation and streptavidin enrichment

For proteomics experiments, all immuno-precipitation steps were performed using an AssayMap Bravo (Agilent) liquid handler in affinity purification v3.0 mode. For antibody loading, 1µg of Anti-FLAG antibodies (Cell Signaling, #2368S) diluted in 100µL PBS were conjugated onto AssayMap 5µL protein A (PA-W) coated cartridges (Agilent, #G5496-60000). Aspiration into the cartridge was at a rate of 4 µg/minute. Cartridges were washed with 50µL 1X PBS at a flow rate of 4µL/minute. For middle-down workflow samples, the entire 255 µL TEV cleavage reaction was loaded directly onto the antibody-conjugated cartridges at an aspiration rate of 4µL/minute. For bottom-up workflow, 100µg of lysate per sample was diluted in lysis buffer to a final volume of 100µL and loaded onto cartridges in the same manner. Cartridges were washed twice with 50µL lysis buffer (1% Triton X-100, 25mM Tris, pH7.5/150mM NaCl, 1mM EDTA/1mM EGTA, UltraPure distilled water) and then twice with 50µL of 1X PBS, both with a flow rate of 4µL/minute. The FLAG-tagged proteins or peptides were eluted with 50 µL of 0.1% Trifluoroacetic acid (TFA) in water (Honeywell, LC485-1) at a flow rate of 2.5µL/minute into a 96-well PCR plate (ABgene SuperPlate, Thermofisher Scientific, #AB-2800). Plates were dried down using a GeneVac for subsequent reduction and alkylation. For experiments treating cells with synthetic biotinylated GCGR agonist and antagonists to enrich active/inactive GCGR (supplemental Figure 1), no quantification of lysate was performed and 250µl of lysate from each sample was loaded directly onto streptavidin-coated AssayMap Bravo 5µl cartridges (Agilent, #G5496-60010). In these experiments the loading, washing, and elution conditions on the AssayMap Bravo were carried out the same as described above for bottom-up experiments except for increased loading volume (250µl vs. 100µl).

#### Reduction, alkylation, tryptic digestion (in 96-well plate)

Following the immuno-precipitation step, both bottom-up and middle-down samples were reduced using 25µL of 25mM ammonium bicarbonate (Sigma-Aldrich, A6141-25g) in water containing 10mM dithiotreitol (DTT, GoldBio, DTT25) and placed at 37°C for 30 minutes (plate sealed prior to incubation). For alkylation, 25 µL of 50mM iodoacetamide (IAA) in 25mM ammonium bicarbonate was added to each well. The plate was then sealed and placed in the dark at room temperature for 30 minutes. At this stage, bottom-up samples were subject to tryptic digestion by adding 5 µL of 100 µg/mL trypsin/Lys-C protease mix (mass spec grade, ThermoFisher Scientific, #A4009) to each sample. The plate was sealed and incubated overnight at 37°C. The next day, the tryptic digest reactions were quenched by the addition of 10 µL of 0.1% TFA (Honeywell, LC485-1). All samples (both TEV enzyme- and trypsin-digested) were filtered using Ultrafree-MC-HV centrifugal filters (Durapore, #UFC30HV00) and returned to a fresh PCR plate (ABgene SuperPlate 96 well, ThermoFisher Scientific, AB-2800) for subsequent proteomic analysis.

#### DDA- and DIA-pasef LC-MS/MS analysis (for bottom-up proteomics workflow)

The bottom-up LC-MS/MS was performed with a NanoElute (Bruker Daltonics) HPLC coupled to a Bruker timsTOF Pro mass spectrometer (Bruker Daltonics) via a nanoelectrospray ion source (Captive Spray, Bruker Daltonics). Two mobile phase solvent systems were utilized for the liquid chromatography (mobile phase A (MPA): 0.1% FA in water; mobile phase B (MPB): 0.1% formic acid in acetonitrile. Each 5 ul of digested sample was loaded on a PepMap Neo C18 300 μm X 5 mm, 5 μm Trap Cartridge (Thermo Scientific, #174500) and separated on a PepSep C18 25 cm × 75 μm, 1.9 μm reversed-phase column (Bruker Daltonics, #1893477) in an oven compartment heated to 40 °C at a flow rate of 350 nL/min using a stepwise mobile phase 80 minutes gradient, from 2% to 25% MPB for 60 minutes, next from 25% to 35% MPB for 10 minutes, then from 35% to 95% MPB for 2 minutes, and finally keeping at 95% MPB for the next 10 minutes. Captive spray capillary was set to 1600V with a drying temperature of 180 °C and gas flow rate of 3 l/min.

For the DDA-PASEF experiments, the instrument was operated with a 1.1 second cycle time DDA-PASEF method comprised of ten 100ms PASEF ramps covering a 1/K0 range between 0.6 Vs/cm^2^ and 1.6 Vs/cm^2^. MS1 scan range was 100 m/z to 1700 m/z with a precursor target intensity of 20,000 and an intensity threshold of 2500. The PASEF precursor ion region was designed to exclude the selection of [M+H] ^+^ precursor ions. Precursor isolation width was 2.00 Th at 700 m/z and 3.0 Th at 800 m/z. Collision energy was set proportionally to 1/k0 with 20.00 V at 0.6 Vs/cm^2^ and 59V at 1.6 Vs/cm^2^.

For the DIA-PASEF experiments, the instrument was operated with an approximately 1.8 second cycle time DIA-PASEF method comprised of 32 DIA isolation events between 400 m/z and 1200 m/z (covering a mobility range between 0.6 Vs/cm^2^ and 1.6 Vs/cm^2^). Each isolation window had a width of 26 Th with a 1 Th overlap. MS1 scan range was between 100 m/z and 1700 m/z. Collision energy was set proportionally to 1/k0 with 20.00 V at 0.6 Vs/cm^2^ and 59V at 1.6 Vs/cm^2^.

#### LC-PRM analysis (for middle-down proteomics workflow)

The targeted middle-down LC-MS/MS was performed with a Vanquish Neo (Thermo Scientific) HPLC coupled to a Thermo Orbitrap Exploris 480 mass spectrometer (Thermo Scientific). Two mobile phase solvent systems were utilized for the liquid chromatography (mobile phase A (MPA): 0.1% FA in water; mobile phase B (MPB): 0.1% formic acid in acetonitrile. Each 5 ul of sample was loaded on a PepMap Neo C18 300 μm X 5 mm, 5 μm Trap Cartridge (Thermo Scientific, #174500) and separated on an EASY-Spray C18 25 cm × 75 μm, 2 μm reversed-phase column (Thermo Scientific, #ES902) heated to 50 °C at a flow rate of 350 nL/min using a stepwise mobile phase 30 minutes gradient, from 1% to 35% MPB for 23.5 minutes, then from 35% to 80% MPB for 0.5 minute, and finally keeping at 80% MPB for the next 6 minutes. For ESI, the spray voltage was set to 1800 V and the ion transfer tube was held at 280°C. The Orbitrap Exploris 480 was operated in a targeted PRM mode (without multiplexing). Precursor ion isolation width was set to 2.0 m/z, HCD collision energies (normalized) for each scan were 30,35,40 (%) with 1 micro-scan. The RF lens % was 50 and the AGC target was set to “Standard” with the orbitrap resolution at 60,000. Data were stored in profile mode. Precursor isolation m/z values for each receptor are listed below.

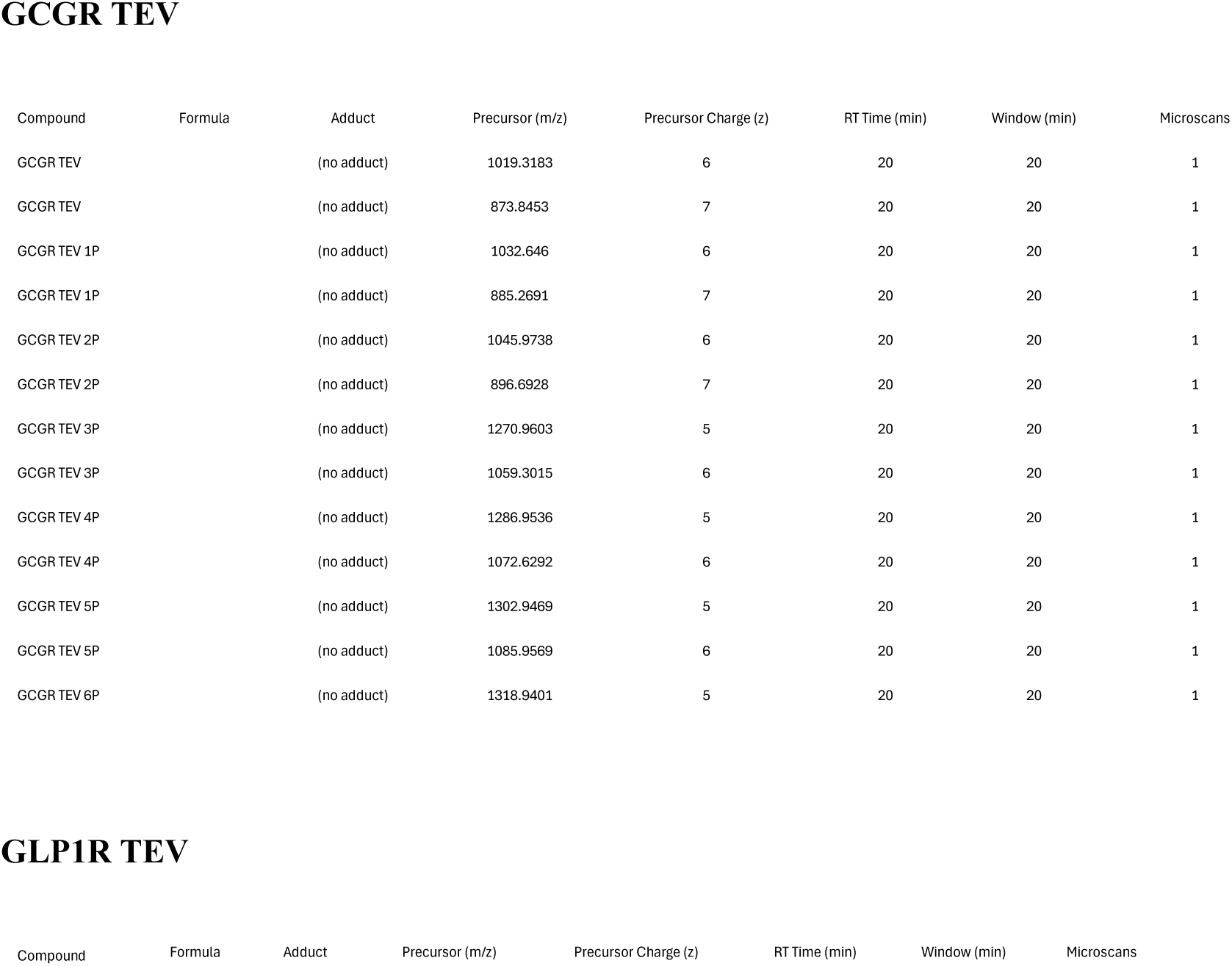
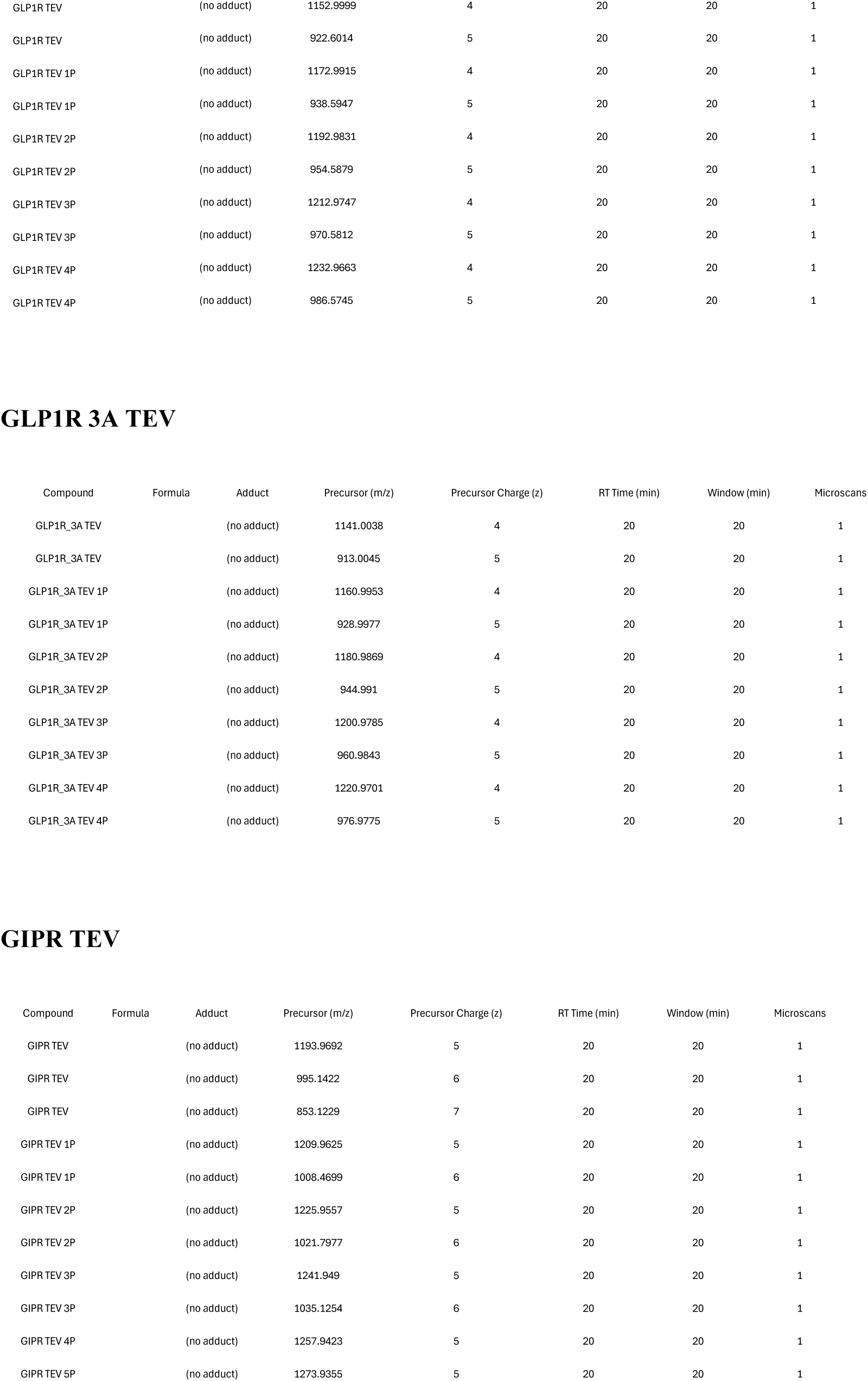

GCGR TEV
| Compound | Formula | Adduct | Precursor (m/z) | Precursor Charge (z) | RT Time (min) | Window (min) | Microscans |
| --- | --- | --- | --- | --- | --- | --- | --- |
| GCGR TEV |  | (no adduct) | 1019.3183 | 6 | 20 | 20 | 1 |
| GCGR TEV |  | (no adduct) | 873.8453 | 7 | 20 | 20 | 1 |
| GCGR TEV 1P |  | (no adduct) | 1032.646 | 6 | 20 | 20 | 1 |
| GCGR TEV 1P |  | (no adduct) | 885.2691 | 7 | 20 | 20 | 1 |
| GCGR TEV 2P |  | (no adduct) | 1045.9738 | 6 | 20 | 20 | 1 |
| GCGR TEV 2P |  | (no adduct) | 896.6928 | 7 | 20 | 20 | 1 |
| GCGR TEV 3P |  | (no adduct) | 1270.9603 | 5 | 20 | 20 | 1 |
| GCGR TEV 3P |  | (no adduct) | 1059.3015 | 6 | 20 | 20 | 1 |
| GCGR TEV 4P |  | (no adduct) | 1286.9536 | 5 | 20 | 20 | 1 |
| GCGR TEV 4P |  | (no adduct) | 1072.6292 | 6 | 20 | 20 | 1 |
| GCGR TEV 5P |  | (no adduct) | 1302.9469 | 5 | 20 | 20 | 1 |
| GCGR TEV 5P |  | (no adduct) | 1085.9569 | 6 | 20 | 20 | 1 |
| GCGR TEV 6P |  | (no adduct) | 1318.9401 | 5 | 20 | 20 | 1 |

GLP1R TEV
| Compound | Formula | Adduct | Precursor (m/z) | Precursor Charge (z) | RT Time (min) | Window (min) | Microscans |
| --- | --- | --- | --- | --- | --- | --- | --- |

|  |  |  |  |  |  |  |
| --- | --- | --- | --- | --- | --- | --- |
| GLP1R TEV | (no adduct) | 1152.9999 | 4 | 20 | 20 | 1 |
| GLP1R TEV | (no adduct) | 922.6014 | 5 | 20 | 20 | 1 |
| GLP1R TEV 1P | (no adduct) | 1172.9915 | 4 | 20 | 20 | 1 |
| GLP1R TEV 1P | (no adduct) | 938.5947 | 5 | 20 | 20 | 1 |
| GLP1R TEV 2P | (no adduct) | 1192.9831 | 4 | 20 | 20 | 1 |
| GLP1R TEV 2P | (no adduct) | 954.5879 | 5 | 20 | 20 | 1 |
| GLP1R TEV 3P | (no adduct) | 1212.9747 | 4 | 20 | 20 | 1 |
| GLP1R TEV 3P | (no adduct) | 970.5812 | 5 | 20 | 20 | 1 |
| GLP1R TEV 4P | (no adduct) | 1232.9663 | 4 | 20 | 20 | 1 |
| GLP1R TEV 4P | (no adduct) | 986.5745 | 5 | 20 | 20 | 1 |

GLP1R 3A TEV
| Compound | Formula | Adduct | Precursor (m/z) | Precursor Charge (z) | RT Time (min) | Window (min) | Microscans |
| --- | --- | --- | --- | --- | --- | --- | --- |
| GLP1R_3A TEV |  | (no adduct) | 1141.0038 | 4 | 20 | 20 | 1 |
| GLP1R_3A TEV |  | (no adduct) | 913.0045 | 5 | 20 | 20 | 1 |
| GLP1R_3A TEV 1P |  | (no adduct) | 1160.9953 | 4 | 20 | 20 | 1 |
| GLP1R_3A TEV 1P |  | (no adduct) | 928.9977 | 5 | 20 | 20 | 1 |
| GLP1R_3A TEV 2P |  | (no adduct) | 1180.9869 | 4 | 20 | 20 | 1 |
| GLP1R_3A TEV 2P |  | (no adduct) | 944.991 | 5 | 20 | 20 | 1 |
| GLP1R_3A TEV 3P |  | (no adduct) | 1200.9785 | 4 | 20 | 20 | 1 |
| GLP1R_3A TEV 3P |  | (no adduct) | 960.9843 | 5 | 20 | 20 | 1 |
| GLP1R_3A TEV 4P |  | (no adduct) | 1220.9701 | 4 | 20 | 20 | 1 |
| GLP1R_3A TEV 4P |  | (no adduct) | 976.9775 | 5 | 20 | 20 | 1 |

GIPR TEV
| Compound | Formula | Adduct | Precursor (m/z) | Precursor Charge (z) | RT Time (min) | Window (min) | Microscans |
| --- | --- | --- | --- | --- | --- | --- | --- |
| GIPR TEV |  | (no adduct) | 1193.9692 | 5 | 20 | 20 | 1 |
| GIPR TEV |  | (no adduct) | 995.1422 | 6 | 20 | 20 | 1 |
| GIPR TEV |  | (no adduct) | 853.1229 | 7 | 20 | 20 | 1 |
| GIPR TEV 1P |  | (no adduct) | 1209.9625 | 5 | 20 | 20 | 1 |
| GIPR TEV 1P |  | (no adduct) | 1008.4699 | 6 | 20 | 20 | 1 |
| GIPR TEV 2P |  | (no adduct) | 1225.9557 | 5 | 20 | 20 | 1 |
| GIPR TEV 2P |  | (no adduct) | 1021.7977 | 6 | 20 | 20 | 1 |
| GIPR TEV 3P |  | (no adduct) | 1241.949 | 5 | 20 | 20 | 1 |
| GIPR TEV 3P |  | (no adduct) | 1035.1254 | 6 | 20 | 20 | 1 |
| GIPR TEV 4P |  | (no adduct) | 1257.9423 | 5 | 20 | 20 | 1 |
| GIPR TEV 5P |  | (no adduct) | 1273.9355 | 5 | 20 | 20 | 1 |

#### Data analysis—proteomic data

The DDA data was searched against homo sapiens (human) proteins database (FASTA format, UniProt version January 2023) which was modified in-house to include C-terminal FLAG-tagged GLP-1R, GCGR and GIPR using Mascot Daemon 2.8.0 (Matrix Science). All searches were performed with carbamidomethyl (C) as a fixed modification, oxidation (M) and phosphorylation (STY) as variable modifications. The search data was analyzed with Scaffold 5.3.3 (Proteome Software) to identify the phosphorylated tryptic peptides.

The DIA data and PRM data was processed with Skyline (64-bit) 24.1.0.199 (MaCoss Lab. Department of Genome Sciences, University of Washington). The target peptides were manually entered into Skyline and the product ions to quantify the intensity (peak area) for every peptide were manually selected, verified and integrated.

The fold change and phosphorylation percentage were calculated using Microsoft Excel. The statistical analysis and Figures were generated with Graph Prism 10.1.2.

#### β-arrestin recruitment assays (BRET)

β-arrestin recruitment to GLP-1R, GCGR, or GIPR was assessed via BRET in HEK Freestyle cells using the NanoBRET detection system (Promega, #N1663). Briefly, cells seeded in suspension at a density of 250,000 cells/mL were transfected with β-arr2 N-terminally fused to NanoLuc and either GLP1R-, GCGR-, or GIPR-Halotag using Fugene-6. After 24 hours, cells were reseeded at a density of 200,000 cells/mL and treated with either 1,000X Halo ligand or DMSO. The next day, cells were pelleted and resuspended in assay buffer (DMEM, #Gibco 30153) containing 0.1% casein and luciferase substrate. Cells were then transferred to 96-well white microplates (Corning, #3917) containing diluted agonists that were prepared via acoustic direct dilution as was done for the aforementioned cAMP accumulation experiments (total reaction volume of 100 µL). Emission was measured at 460 nm for donor, as well as 610 nm for the acceptor, wavelengths using an EnVision plate reader, and signals were acquired for approximately 10-15 minutes until a plateau was achieved. Data analysis was performed using GraphPad Prism 10.1.2.

#### cAMP accumulation assays (HTRF)

HEK Freestyle cells (ThermoFisher Scientific, #R79007) seeded in suspension at a density of 250,000 cells/mL were transiently transfected with indicated receptor constructs using Fugene-6 reagent (Promega, #E2691). After 48 hours, cells were pelleted and resuspended in assay buffer (DMEM, #Gibco 31053) containing 0.1% casein. Cells were then added at 1000 cells/well to 384-well white microplates (Costar, #3570) containing a range of agonist concentrations prepared via acoustic direct dilution in assay buffer containing 250 µM IBMX (total reaction volume of 20 µL), followed by incubation at 37 °C for 30 minutes. Cells were then lysed via sequential addition of d2-labeled cAMP competitor conjugate and cryptate-conjugated detection antibody (Revity, #62AM4PEC), then incubated for 1 hour at room temperature with subsequent quantification of time-resolved fluorescence resonance energy transfer using an Envision plate reader and calibration to external synthetic cAMP standards in a parallel processed plate. Normalized percent values were fit to the 4-parameter logistic model using GraphPad Prism 10.1.2.

## Results

We examined ligand-induced phosphorylation of the GCGR transiently expressed in HEK293 cells using a bottom-up proteomic approach. Cells transiently expressing GCGR-FLAG were stimulated with 1 µM GCG (1-29) or DMSO for 10 minutes, followed by immunoprecipitation of the receptor and enzymatic digestion for subsequent LC-MS/MS analysis (Figure 1A). We observed three serine residues that exhibited ligand-dependent increases in phosphorylation relative to DMSO treatment, namely Ser445, Ser456, and Ser459 (Figure 1B-C). We also observed a slight but significant decrease in Ser438 phosphorylation upon the addition of ligand (Figure 1B). In contrast, Ser475 appeared to be phosphorylated under basal conditions and was not further modified upon the addition of agonist (Figure 1D). Importantly, the quantities of the associated unmodified peptides identified in our LC-MS/MS analysis were not significantly changed in agonist-treated samples compared to DMSO-treated samples, suggesting these phosphorylation changes were not due to relative peptide abundance between the samples (Figure 1B-D). A summary diagram of the locations of these PTMs in the GCGR is shown in Figure 1E. To assess the functional significance of these observed PTMs, we mutated each of these residues to alanine and measured agonist stimulated cAMP accumulation and βarr recruitment to the GCGR using Homogenous Time Resolved Fluorescence (HTRF) and Bioluminescence Resonance Energy Transfer (BRET) assays, respectively. Strikingly, simultaneous mutagenesis of all five phosphorylated C-tail residues to alanine (“5S/A”) showed no modulation of either cAMP production or βarr recruitment (Figure 1F-G). Additionally, individual single mutants S445A, S456A, S459A, nor a combination mutant of all three of these residues (“3S/A”), conferred modulation of cAMP production or βarr recruitment (Figure 1F-G).

**Figure 1.**
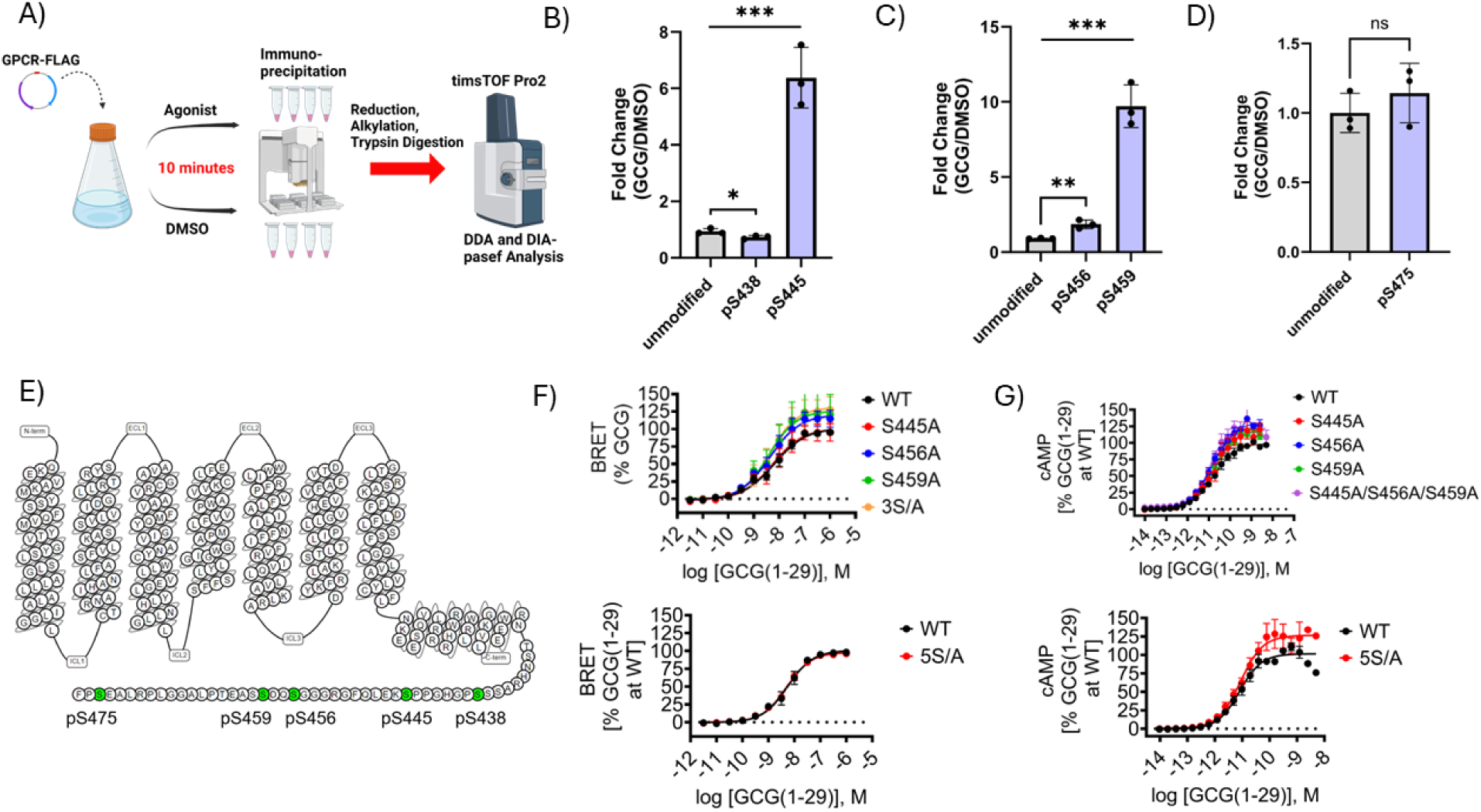
Detection of C-tail phosphorylation and mutational analysis of GCGR signaling properties. A) Schematic diagram of bottom-up proteomic method used to identify phosphorylated residues in GCGR C-tail. B-D) Quantification of fold change of phosphorylation in GCGR C-tail residues in cells treated with 1 µM GCG (1-29) compared to DMSO. E) Topological map of the GCGR with phosphorylated C-tail residues are shown in green. F) BRET assay measuring nano-luciferase fused β-arr2 (nLuc-β-arr2) recruitment to WT GCGR-Halo receptor compared to individual S445A, S456A, S459A mutants, a combination triple mutant of these three residues (3S/A), and a 5S/A mutant in which all five residues we detected to be phosphorylated were mutated to alanine (pS438/pS445/pS456/pS459/pS475). G) cAMP accumulation assay of WT GCGR-FLAG receptor compared to the same mutants tested in F. For proteomics experiments, data points and error bars are the mean and SD of averaged technical quadruplicates from three independent experiments. An unpaired two-tailed t test was conducted for statistical analysis; n=3. ns, nonsignificant; *\*P* < 0.05; *\*\*P* < 0.01; *\*\*\*P* < 0.001. For pharmacology experiments summary statistics are presented in Table S1.

To unambiguously define the phosphorylation state of the activated GCGR, we employed an orthogonal approach utilizing biotinylated peptide ligands to enrich GCGR for subsequent LC-MS/MS analysis. Biotinylated GCG (1-29)-derived GCGR agonist and antagonist peptides were generated as affinity reagents to facilitate the enrichment of the active and inactive receptor species (supplemental Figure 1C). Importantly, these biotinylated peptides retained pharmacological properties consistent with non-biotinylated analogues (supplemental Figure 1A-B). In agreement with our initial findings, phosphorylation of Ser445 was again increased in agonist-treated samples (supplemental Figure 1D). Furthermore, Ser438 again displayed modestly decreased phosphorylation in biotinylated agonist-treated samples relative to antagonist in these experiments (supplemental Figure 1D). In contrast to our initial set of experiments, Ser456 had no change in magnitude of phosphorylation in agonist-treated samples relative to antagonist (supplemental Figure 1E), and Ser459/Ser475 were not identified using this method. Overall, the results of these experiments largely confirmed the identity of key C-tail residues that undergo modulation of phosphorylation in GCGR following agonism and provided confidence in our initial results such that this workflow could be successfully applied to the other glucagon family receptors. Critically, the unexpected findings regarding lack of effects of Ser to Ala mutation of GCGR phosphorylated C-tail residues on signaling led us to question whether the glucagon family of receptors are reliant upon C-tail phosphorylation for proper function.

To determine the GIPR residues phosphorylated upon activation, we applied the same bottom-up proteomic workflow as we did for the GCGR (shown in Figure 1A). With respect to C-tail phosphorylation, we detected significant GIP (1-42)-induced phosphorylation at three putative locations: Ser433 or S435, Ser443, and Ser447 or Ser448 (Figure 2A, Figure 2C). In the case of Ser433/Ser435 and Ser447/Ser448 residue pairs, we were unable to determine the precise location of phosphorylation in the C-tail region due to a lack of sequence-specific product ions in our MS/MS data. We also observed a GIP (1-42)-mediated enhancement of phosphorylation in the helix 8 domain of GIPR at Ser415 (Figure 2B). Unexpectedly, we also detected a phosphorylated residue in the transmembrane 6 domain of GIPR at Ser342 or Tyr343 (Figure 2D). This residue appears to be basally phosphorylated and was not modulated upon GIPR activation (Figure 2D). The quantities of unmodified peptide species associated with these PTMs were equal in GIP-treated samples compared to DMSO-treated samples, suggesting these phosphorylation changes were not due to relative peptide abundance between the samples (Figure 2A-D). In contrast to our observations for GCGR, mutation of individual phosphorylated residues in GIPR were sufficient to modulate signaling. For example, individual S415A, S433A, S447A, and S448A mutants all caused a decrease in βarr recruitment relative to wild type receptor (Figure 2F), as would be expected under the classical model. These mutants also displayed an increase in cAMP accumulation (Figure 2G).

**Figure 2.**
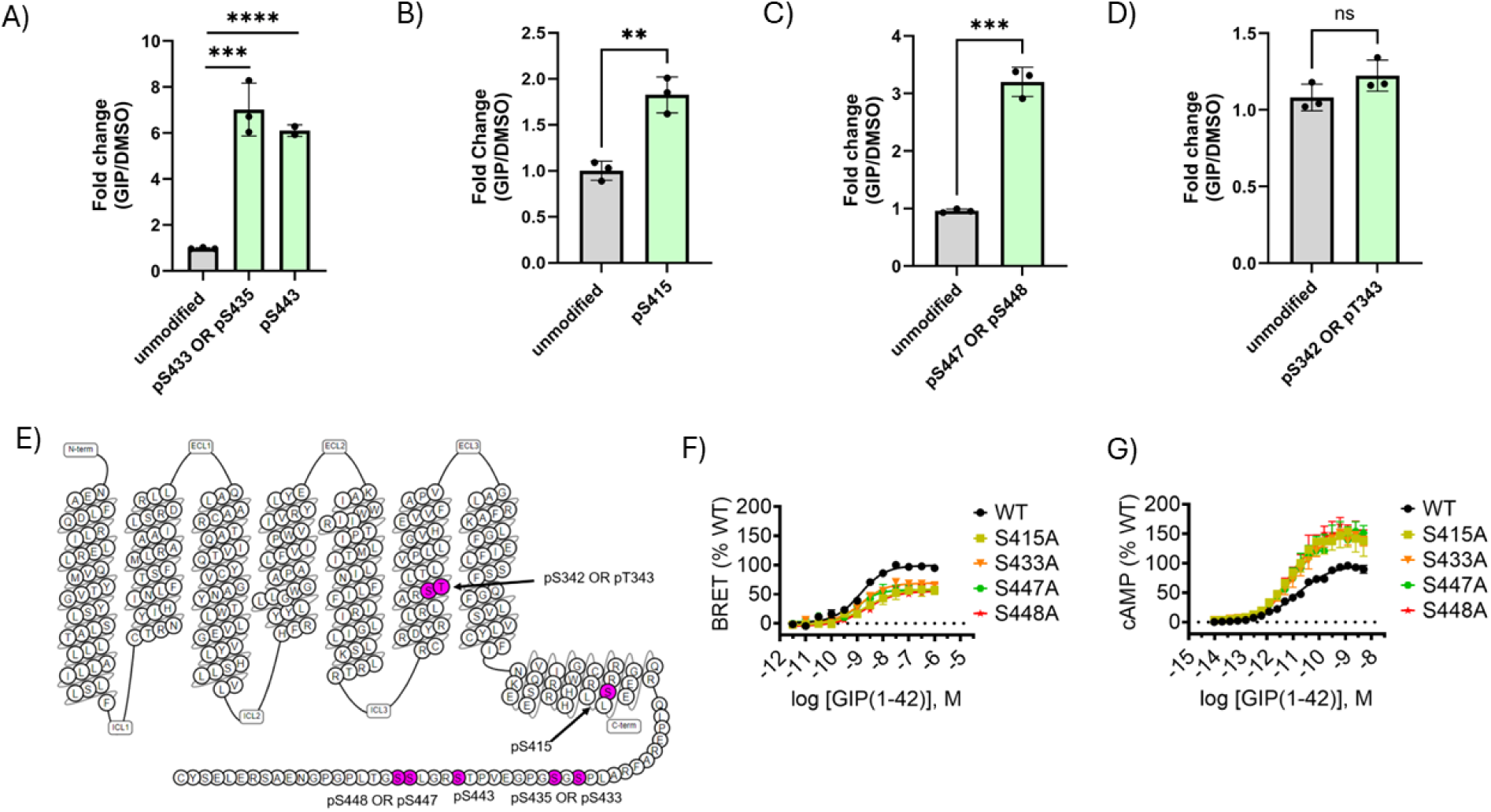
Bottom-up proteomic detection of C-tail phosphorylation and mutational analysis of GIPR signaling properties. A-D) Quantification of fold change of phosphorylation in GIPR residues in cells treated with 1µM GIP (1-42) compared to DMSO. E) Topological diagram of GIPR with phosphorylated residues shown in pink. F) BRET assay measuring nano-luciferase fused β-arr2 (nLuc-β-arr2) recruitment to WT GIPR-Halo tagged receptor compared to individual S415A, S433A, S447A, and S448A mutants. G) cAMP accumulation assay of WT GIPR-FLAG receptor compared to the same mutants tested in F. For proteomics experiments, data points and error bars are the mean and SD of averaged technical quadruplicates from three independent experiments. An unpaired two-tailed t test was conducted for statistical analysis; n=3. ns, nonsignificant; *\*\*P* < 0.01; *\*\*\*P* < 0.001.; *\*\*\*\*P* < 0.0001. For pharmacology experiments summary statistics are presented in Table S1.

Finally, we wanted to determine which C-tail residues are phosphorylated in the third glucagon receptor family member, GLP-1R, in response to agonism and characterize the effects of mutagenesis of these residues on receptor signaling. We applied a bottom-up proteomic workflow analogous to that used for GCGR and GIPR (shown in Figure 1A). Despite obtaining good C-tail coverage in the LC-MS/MS data for GLP-1R (supplemental Figure 2), we did not detect any phosphorylation. Subsequent experiments with Immobilized Metal Affinity Chromatography (IMAC) enrichment of the tryptic digest following immunoprecipitation of the receptor also failed to identify any phospho-peptides in GLP-1R (data not shown). To obviate this, we implemented a recently established proteomic method to identify C-tail phosphorylation in GPCRs ^19^. This “middle-down” approach makes use of the insertion of a Tobacco Etch Virus (TEV) recognition sequence directly after transmembrane domain seven and the addition of an affinity purification tag at the C-terminus of the receptor. Following TEV cleavage of the receptor, the entire intact C-tail is enriched by immuno-precipitation and analyzed by Liquid Chromatography-Parallel Reaction Monitoring (LC-PRM) (Figure 3A). This targeted middle-down method increases the depth of C-tail coverage and enhances the detection of low-abundance PTMs relative to the “bottom-up” method shown in Figure 1A, which analyzes tryptic peptides generated from the entire receptor. Compared to bottom-up approaches in which co-occurring modifications are separated by peptide cleavage, this method provides increased quantitative information regarding the number of co-occurring phosphorylated residues (stoichiometry) on each receptor and relative abundances of each modification state or, “proteoform”. Employing this technique, we generated a GLP-1R construct in which the TEV recognition sequence was inserted between Ser431 and Ser432 at the junction between helix 8 and the C-tail, and a FLAG epitope tag was installed at the C-terminus. We first confirmed that insertion of the TEV recognition sequence did not significantly alter βarr recruitment to the receptor following GLP-1 (7-36) treatment (data not shown). We then used this construct to carry out our middle-down workflow (Figure 3A) to quantify ligand-dependent phosphorylation in the GLP-1R C-tail following treatment of cells with either 1 µM of GLP-1 (7-36) (Figure 3B) or the small molecule agonist danuglipron (Figure 3C) relative to DMSO ^22^. We observed a 14% or 14.5% increase in the percentage of phosphorylated GLP-1R proteoforms after GLP-1 or danuglipron treatment, respectively (Figure 3B-C). In the chromatograms generated in these experiments, we noticed that phosphorylated GLP-1R C-tail exists in distinct and diverse low-abundance mono-, di-, and tri-phosphorylated proteoforms (Proteoforms 1A/1B/1C/1D, 2A/2B, 3A/3B, respectively) following agonism (supplemental Figure 3 B-D) in addition to the unmodified proteoform (0P, supplemental Figure 3A). These species were not identified using the bottom-up proteomic method. We were able to assign proteoform 1B to phospho-Ser442 (pSer442), proteoform 1C to either pSer444 or pSer445, and proteoform 2B to pSer442 and pSer444 OR pSer442 and pSer445 using the *de novo* sequencing of the MS/MS spectra (supplemental Figure 4). One of the 3P proteoforms are thus presumably attributable to pSer442, pSer444, and pSer445; our *de novo* sequencing results suggest that the other is a combination of two of these serine residues along with pSer441. We were not able to assign the precise identities of proteoforms 1A/1D/2A/3A/3B. Figure 3F shows a toplogical diagram of the location of these amino acids on the GLP-1R. Proteoform 1B (Ser442), 2B (Ser442 and Ser444 OR Ser442 and Ser445), and 3A/3B underwent marked increases in phosphorylation after treatment with both GLP-1 and danuglipron (Figure 3D-E). Encouragingly, the proteoform profiles look nearly identical after stimulation with either GLP-1 or danuglipron agonists (Figure 3D-E), which interestingly have similar signaling properties but bind at different locations on the receptor. To determine the functional impact of these ligand-induced modifications on receptor signaling, we generated a mutant construct for GLP-1R in which Ser442, Ser444, and Ser445 were mutated either individually or as a cluster (S442A/S444A/S445A) to alanine. We assessed β-arr recruitment and cAMP accumulation of these mutants upon treatment with both GLP-1 and danuglipron (Figure 3G-H). With both ligands, we observed only a minor decrease in β-arr recruitment to the GLP-1R in the single mutants (S442A or S444A or S445A) (Figure 3G). Simultaneous mutation of all three of these residues was required for a significant defect in β-arr recruitment. We did not detect a significant difference in cAMP accumulation between the individual S442A/S444A/S445A or cluster mutants and wild type GLP-1R (Figure 3H).

**Figure 3.**
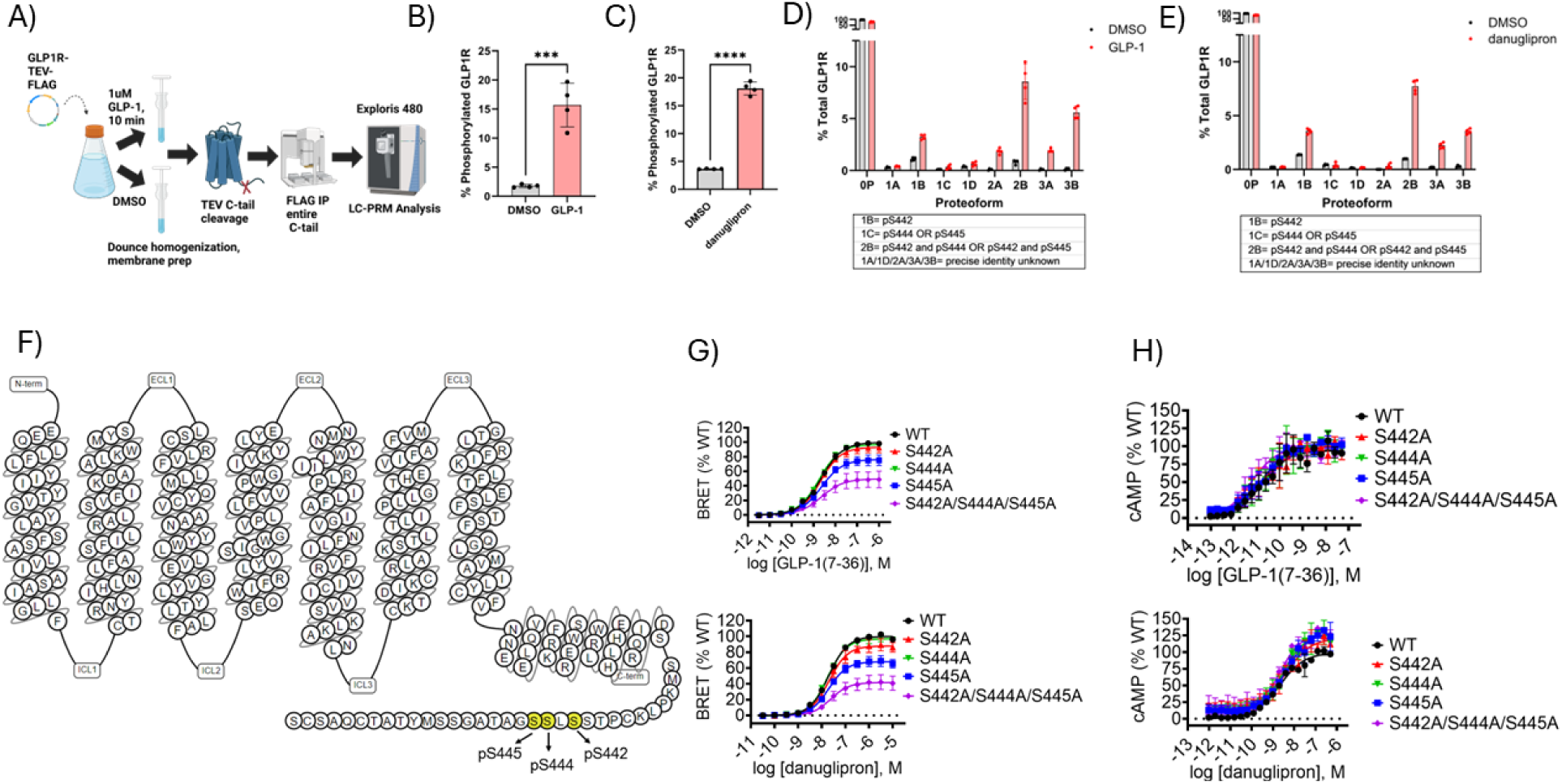
Detection of C-tail phosphorylation and mutational analysis of GLP-1R signaling properties. A) Schematic diagram of middle-down proteomic workflow used to identify phosphorylated residues in the GLP-1R_TEV_FLAG C-tail. B-C) Bar graphs indicating the percentage of GLP-1R C-tail that is phosphorylated following treatment with 1µM GLP-1(7-36) (B) or danuglipron (C). An unpaired two-tailed t test was conducted for statistical analysis. Data points and error bars are the mean and SD of technical quadruplicates from a representative example of three independent experiments; *\*\*\*P* < 0.001.; *\*\*\*\*P* < 0.0001. D) Bar graph indicating the percentage of each discrete proteoform among total GLP-1R after treatment with DMSO (grey bars) or GLP-1(7-36) (red bars). Data points and error bars are the mean and SD of technical quadruplicates from a representative example of three independent experiments. 0P is an unmodified proteoform; 1A-1D are mono-phosphorylated proteoforms; 2A-2B are di-phosphorylated proteoforms; 3A-3B are tri-phosphorylated proteoforms. If they could be determined by *de novo* sequencing of MS/MS spectra, the phosphorylated residue(s) associated with these species are shown in a box below the bar graphs. E) Same experimental setup as D but using danuglipron (red bars) as the ligand. Data points and error bars are the mean and SD of technical quadruplicates from a representative example of two independent experiments. F) Snake diagram of GLP-1R showing key residues (in yellow) that undergo modulation of phosphorylation with GLP-1(7-36) treatment relative to DMSO. G) BRET assay measuring nLuc-β-arr2 recruitment to WT GLP-1R-Halo receptor compared to the individual single mutant receptors S442A, S44A, S445A as well as a cluster mutant of all three of these residues in cells treated with GLP-1(7-36) or danuglipron. H) cAMP accumulation assay of WT GLP-1R-FLAG vs. the same mutants as in (G) in cells treated with GLP-1(7-36) or danuglipron. For pharmacology experiments summary statistics are presented in Table S1.

Having identified S442/S444/S445 as key GLP-1R C-tail residues that are phosphorylated in a ligand dependent manner, we wanted to determine the effect of alanine substitution of these residues on C-tail phosphorylation. We thus generated a phospho-deficient mutant S442A/S444A/S445A GLP-1R_TEV receptor with a C-terminal FLAG tag and repeated the middle down proteomic workflow shown in Figure 3A using this construct. We measured only a 1% increase in the percentage of phosphorylated GLP-1R C-tail proteoforms in this experiment (supplemental Figure 5A), a large decrease in modulation of phosphorylation relative to the 14% increase we detected in WT GLP-1R_TEV C-tail proteoforms following GLP-1(7-36) treatment (Figure 3B). This result provides evidence that S442/S444/S445 are indeed the “canonically” phosphorylated residues in GLP-1R following GLP-1 stimulation. Interestingly, in the context of our S442A/S444A/S445A phospho-deficient C-tail mutant of GLP-1R, we measured phosphorylation at “alternate” residues that were not identified in the WT GLP-1R_TEV experiments (proteoforms 1E-1G; 2C-2F; 3C-3E; 4P, supplemental Figure 5B-G). Together with our data from WT GLP-1R_TEV experiments (Figure 3B-E, supplemental Figure 3A-D), these results suggest that there are canonically phosphorylated residues in GLP-1R following GLP-1 treatment, but that GRKs phosphorylate alternate residues when these sites are ablated (supplemental Figure 5). However, the quantity of phosphorylated receptor is much lower at the alternate residues (supplemental Figure 5A-B) compared to the canonically phosphorylated residues (Figure 3B and 3D). We also did not measure a robust ligand-dependent increase in the alternate di- and tri-phosphorylated proteoforms (supplemental Figure 5B) as we did for WT GLP-1R_TEV receptor following GLP-1 stimulation (Figure 3D). Given that we observed a marked decrease in βarr recruitment by BRET with the GLP-1R S442A/S444A/S445A-Halo mutant receptor (Figure 3G), these results suggest that the di- and tri-phospho-proteoforms (combinations of pS442, pS444, pS445, and presumably pS441) are critical to promoting GLP-1R-βarr complex formation.

For GLP-1R, we were unable to detect any C-tail phosphorylation using our bottom-up workflow and obviated this challenge by employing the middle-down proteomic method. In the interest of extending our proteomic analyses to all three receptors using both bottom-up and middle-down workflows, we applied our middle-down method to GCGR and GIPR (Figure 3A). Thus, we created constructs in which a TEV recognition sequence was inserted at the junction between helix 8 and the C-tail for both receptors (GCGR_N429_TEV_T430; GIPR_R421_TEV_Q422) in addition to a C-terminal FLAG tag. Interestingly, GCGR appeared to have a high percentage of basal C-tail phosphorylation (49% of total C-tail proteoforms) in DMSO treated samples (supplemental Figure 6A)—substantially higher than we observed for WT GLP-1R (1.8%, Figure 3B) and GIPR (1.5%, supplemental Figure 6C) using the same method. GCGR also had the highest increase in the percentage of phosphorylated C-tail proteoforms following agonism (21%, supplemental Figure 6A), GIPR had the smallest increase (5%, supplemental Figure 6C), and GLP-1 fell between these receptors with a 14% increase in phosphorylated C-tail proteoforms (Figure 3B). Similar to the WT GLP-1R middle-down results, we measured marked percentage increases in ligand-dependent phosphorylation in the form of multiple discrete C-tail proteoforms for both GCGR (supplemental Figure 6B, supplemental Figure 7A-F) and GIPR (supplemental Figure 6D, supplemental Figure 8A-D). Of note, GCGR had the most phosphorylated C-tail residues (up to penta-phosphorylated, supplemental Figure 6B, supplemental Figure 7A-F) of all three receptors following agonist stimulation, while both WT GLP-1R and GIPR only had up to tri-phosphorylated proteoforms (supplemental Figure 3, supplemental Figure 8, respectively.)

## Discussion

Classically, ligand-induced phosphorylation of multiple residues in the GPCR C-tail is thought to promote physical interaction between βarrs and the receptor ^2, 3^. These interactions lead to conformational changes in both βarrs and the receptor that, in turn, yield complex cellular signaling cascades. However, the governing principles of complex formation between phosphorylated receptor C-tail and βarrs, and how each individual phosphate modification contributes to modulation of βarr structural/functional states and thus signaling, remain elusive. In this study, we used mass spectrometry approaches to identify four C-tail residues in GCGR that were differentially phosphorylated in a ligand-dependent manner (pS438, pS445, pS456, pS459) and one residue that was basally phosphorylated (pS475) (Figure 1B-D). When we mutated all five of these C-tail residues to alanine, unexpectedly, we observed no change in GCGR-βarr proximity as measured by BRET (Figure 1F). This result prompted us to question how βarrs are recruited to the GCGR in this phospho-deficient context. One possibility is that alternate residues are phosphorylated in our 5S/A GCGR mutant following agonism, much like we observed for the S442A/S444A/S445A GLP-1R_TEV triple mutant receptor (supplemental Figure 5), and that this is sufficient to promote interaction with βarrs to the same degree as WT GCGR receptor. Indeed, there are two additional threonine residues and four additional serine residues in our phospho-deficient GCGR C-tail mutant which may fulfill this function (Figure 1E). Previous reports have shown that mutation of specific groups of canonically phosphorylated residues does not eliminate binding of βarrs to GPCRs in a cellular context so long as additional phosphorylated residues are present ^23, 24^. It has thus been suggested that any phosphorylated C-tail peptide can, at least to a degree, stabilize activated βarr2 ^25^. Intriguingly, this does not appear to be the case for GLP-1R: in our phospho-deficient S442A/S44A/S445A triple mutant receptor, we observed a decrease in βarr recruitment relative to WT receptor (Figure 3G). There are six additional serine residues and four additional threonine residues in this mutant GLP-1R C-tail (Figure 3F), but these do not appear to compensate for the loss of pS442/pS444/pS445 with respect to βarr recruitment.

Another possibility to explain our GCGR 5S/A βarr recruitment results is that, in the presence of GCG, negatively charged acidic resides within the GCGR C-tail are sufficient to promote receptor-βarr interaction. Such phosphomimetic ^26^ residues (aspartic acid and glutamic acid) provide a negative charge and similar volume to a phosphate group, and are thus widely used in protein science to mimic phosphorylated residues in proteins for functional studies ^27, 28^. Specifically, phosphomimetics have been used in a recent investigation to promote GPCR-βarr interaction for structural characterization ^29^. Moreover, a recent study reported crystal structures of βarr2 in complex with four synthetic phospho-peptides derived from the C-tail of the vasopressin receptor-2 (V2R) GPCR ^30^. In comparing their structures to published GPCRs in complex with βarr2 ^31–33^, Q.T. He *et al.* produced a model detailing specific phosphorylated or negatively charged (phosphomimetic) V2R C-tail residues (Asp355 and Glu356) and their associated binding pockets in βarr2 ^30^. Determining whether acidic C-tail residues in the phospho-deficient GCGR C-tail contribute to complex formation with βarrs will require additional experimentation. In addition to negatively charged residues promoting receptor-βarr2 interaction, there is also biochemical evidence that the central transmembrane core (TM core) of a class A GPCR, neurotensin receptor 1, (NTSR_1_) is sufficient for complex formation with βarr2, even in the absence of receptor phosphorylation ^34^. Whether the GCGR TM core has a role in facilitating interactions with βarr2, especially in a phospho-deficient C-tail context, will require further investigation. Overall, the results of our work and others suggest that the existence of receptor-specific “phosphorylation barcodes”, their impact on the structure of receptor-bound βarrs and associated downstream signaling events, remains a worthy subject of further examination.

Recently, a bottom-up proteomic approach was used to determine residues in the GIPR that are phosphorylated in response to agonism ^18^. The approach Brown *et al.* used was similar to the bottom-up workflow we carried out to identify ligand-induced phosphorylation in the GCGR and GIPR (Figures 1 and 2, respectively). Encouragingly, our GIPR LC-MS/MS results are concordant with the residues previously reported for GIPR ^18^. Notable differences were that GIPR pS433 was only identified in our study, while pS459 and pS464 were only identified in Brown *et al.* ^18^. In Brown *et al.,* alanine substitution mutants of the phosphorylated GIPR residues were not generated and thus the function of these PTMs regarding receptor signaling was not investigated. Considering this, we wanted to perform mutational analysis of the phosphorylated GIPR residues we identified using cAMP accumulation and βarr recruitment assays. We found that individual mutation of single phosphorylated C-tail residues was sufficient to significantly decrease βarr recruitment to the GIPR (Figure 2F). These results were in stark contrast to our signaling results for GCGR, where single phospho-deficient mutants, a combination triple mutant (3S/A), nor simultaneous mutation of all five phosphorylated C-tail residues that we identified by LC-MS/MS (5S/A) had any effect on βarr recruitment (Figure 1F). GLP-1R seems to fall into yet a third category in this family of receptors—ablation of individual phosphorylated C-tail residues conferred only a minor decrease of βarr recruitment, while simultaneous mutation of the three phosphorylated C-tail residues caused a large defect in βarr recruitment to the receptor (Figure 3G). Overall, it is not evident that a universal mechanism for βarr recruitment by glucagon-family (and ClassB1) receptors exists ^25^. It is likely that detailed structural studies will be required to full understand the recruitment of βarr to these receptors ^25^.

One of the biggest technical challenges we encountered over the course of this investigation was our inability to detect phosphorylated C-tail residues in the GLP-1R C-tail using the bottom-up proteomic method despite detection of peptides covering most of this region ([S432-S463], (supplemental Figure 2)). At the same time, as expected, we detected robust ligand-induced C-tail phosphorylation in both GCGR (Figure 1B-D) and GIPR (Figure 2A-D) using this method. We reasoned that, in contrast to GCGR and GIPR, phosphorylation of the GLP-1R C-tail constitutes very low-abundance modifications that are below the limit of detection or signal-to-noise ratio of our assay. Accordingly, to examine the relative stoichiometry of C-tail phosphorylation between these three receptors, we employed a middle-down proteomic method recently developed to detect C-tail phosphorylation in GPCRs as a percent of total receptor ^19^. In examining the middle-down data for all three receptors, we noticed that GIPR actually measured the lowest percentage of phosphorylated C-tail proteoforms following agonism with endogenous ligand (6.7%, supplemental Figure 6C) while GLP-1R (Figure 3B) and GCGR (supplemental Figure 6A) measured 15.7% and 71.65%, respectively. It is thus unlikely that a relatively low abundance of C-tail phosphorylation in GLP-1R explains our inability to detect it via the bottom-up proteomic approach. One possible explanation is that phosphorylation at the GLP-1R C-tail had a negative impact on the efficiency of tryptic digest, and thus only unmodified C-tail peptides were generated for LC-MS/MS analysis. Indeed, phosphorylation has been shown to decrease the efficiency of tryptic digestion of specific sequences ^35, 36^ Another possibility is that, for unknown reasons, the particular GLP-1R C-tail phospho-peptides we generated in our tryptic digests are recalcitrant to LC-MS/MS analysis due to their specific chemical composition. Overall, this study highlights that proteomic methods are not universally applicable. We propose the integrated application of complimentary approaches, such as bottom-up and middle-down proteomic workflows, are required to definitively analyze PTMs of complex membrane proteins such as GPCRs.

## Data Availability

All mass spectrometry proteomics data and the associated meta-data have been deposited into a publicly available repository at https://repository.jpostdb.org/ (jPOSTrepo). For the middle-down data, the accession numbers are PXD060260 for ProteomeXchange and JPST003581 for jPOST. For the bottom-up data, the accession numbers are PXD060263 for ProteomeXchange and JPST003580 for jPOST. The identities of each raw file with respect to figure number and replicate are available in Supplementary Table 2.

## Acknowledgments

We thank Caitlyn Keck for technical assistance with the proteomics experiments. We also thank Yuewei Qian for help with plasmid generation.

## Author contact information

Ian M. Lamb, Alex D. White, Francis S. Willard, Michael J. Chalmers, Junpeng Xiao.

## Supplemental Information

**Supplemental Figure 1.**
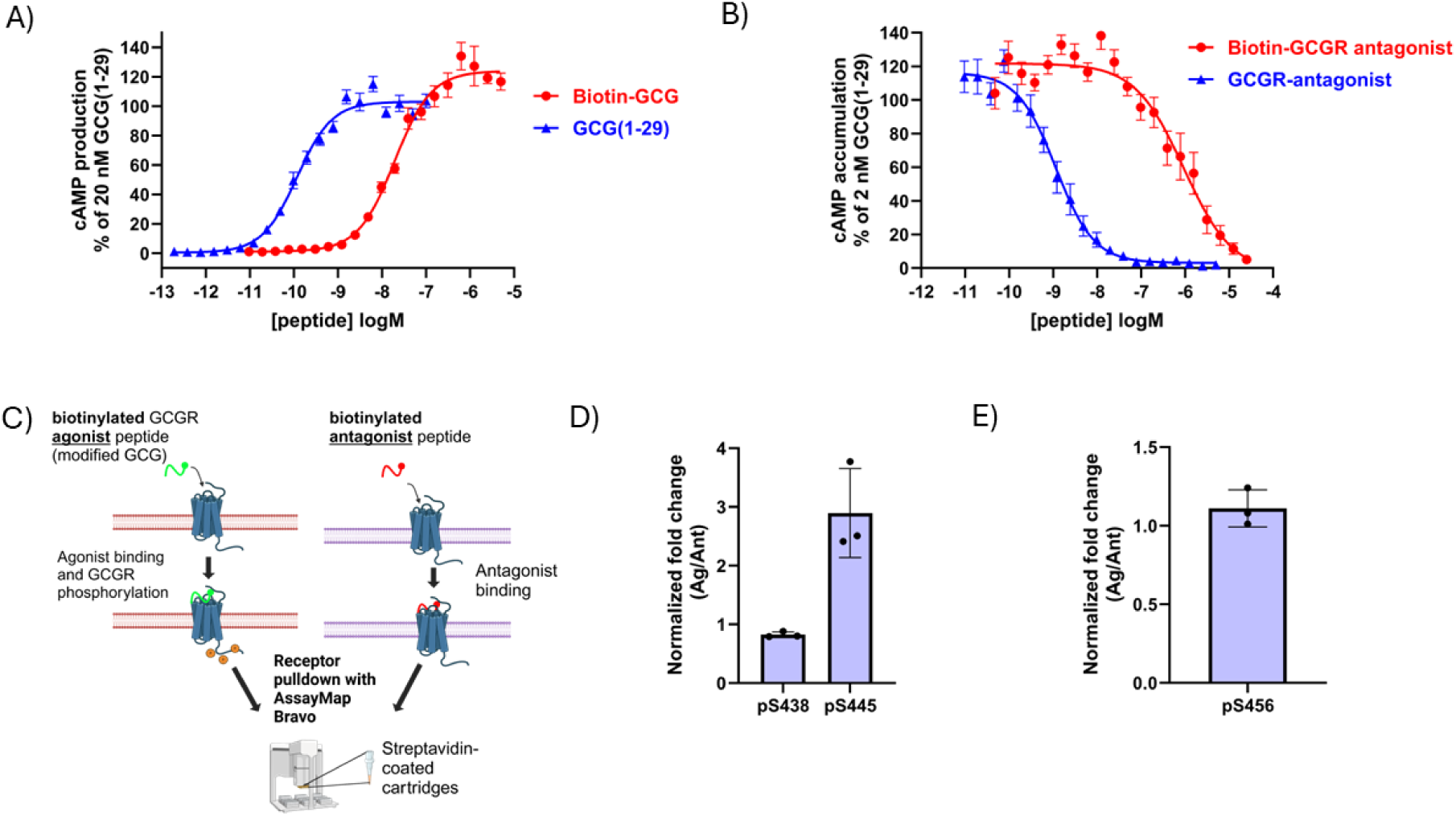
Enrichment of GCGR using biotinylated peptide ligands and bottom-up proteomic analysis of C-tail phosphorylation. A-B) GCGR cAMP accumulation was measured in (A) agonist mode and (B) antagonist mode using native and biotinylated peptides. Summary statistics for pharmacological data are presented in Table S1. C) Schematic diagram of method to enrichment of antagonist-bound GCGR for proteomic analysis. 1 µM of each ligand was used for enrichment. D-E) Quantification of C-tail phosphorylation of agonist-bound receptor relative to antagonist-bound receptor at the three residues detected to be phosphorylated. Normalization was done to account for the efficiency difference of enrichment between biotinylated agonist vs. antagonist. For each peptide containing a phosphorylated residue, normalization was done by summing the peak area for all proteoforms (modified and unmodified) from all samples in each treatment group (agonist vs. antagonist treated) to get a grand total peak area for agonist treated samples and for antagonist treated samples. For each individual sample, the peak area of each proteoform was divided by the agonist or antagonist grand total peak area to obtain the percentage abundance, which was used to calculate the fold change (agonist/antagonist) as a function of percent abundance of that proteoform. Data points and error bars are the mean and SD of averaged technical quadruplicates from three independent experiments.

**Supplemental Figure 2.**
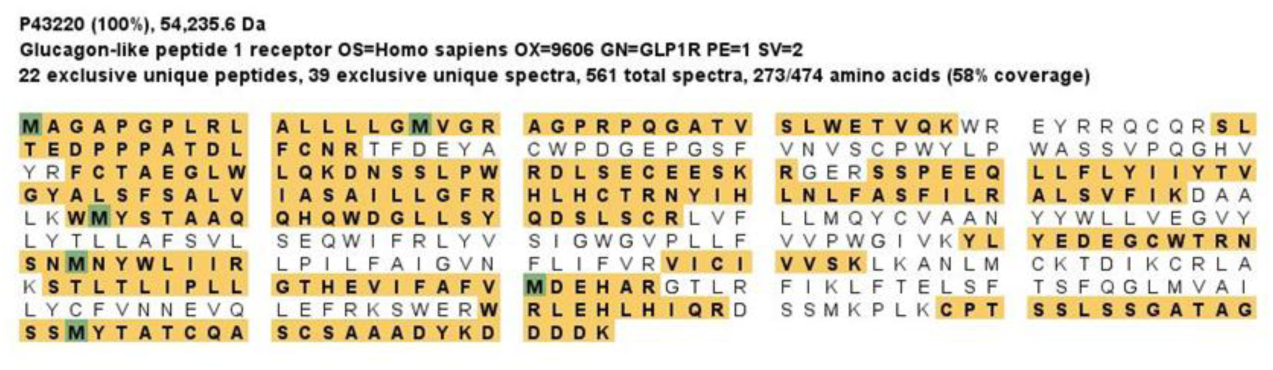
Bottom-up proteomic analysis of GLP-1R C-tail phosphorylation. Graphical display of sequence coverage of GLP-1R-FLAG proteomic analysis. Peptides detected by LC-MS/MS are highlighted in yellow; amino acids bearing a PTM are highlighted in green. Note the absence of serine phosphorylation in the C-tail or anywhere on the receptor.

**Supplemental Figure 3.**
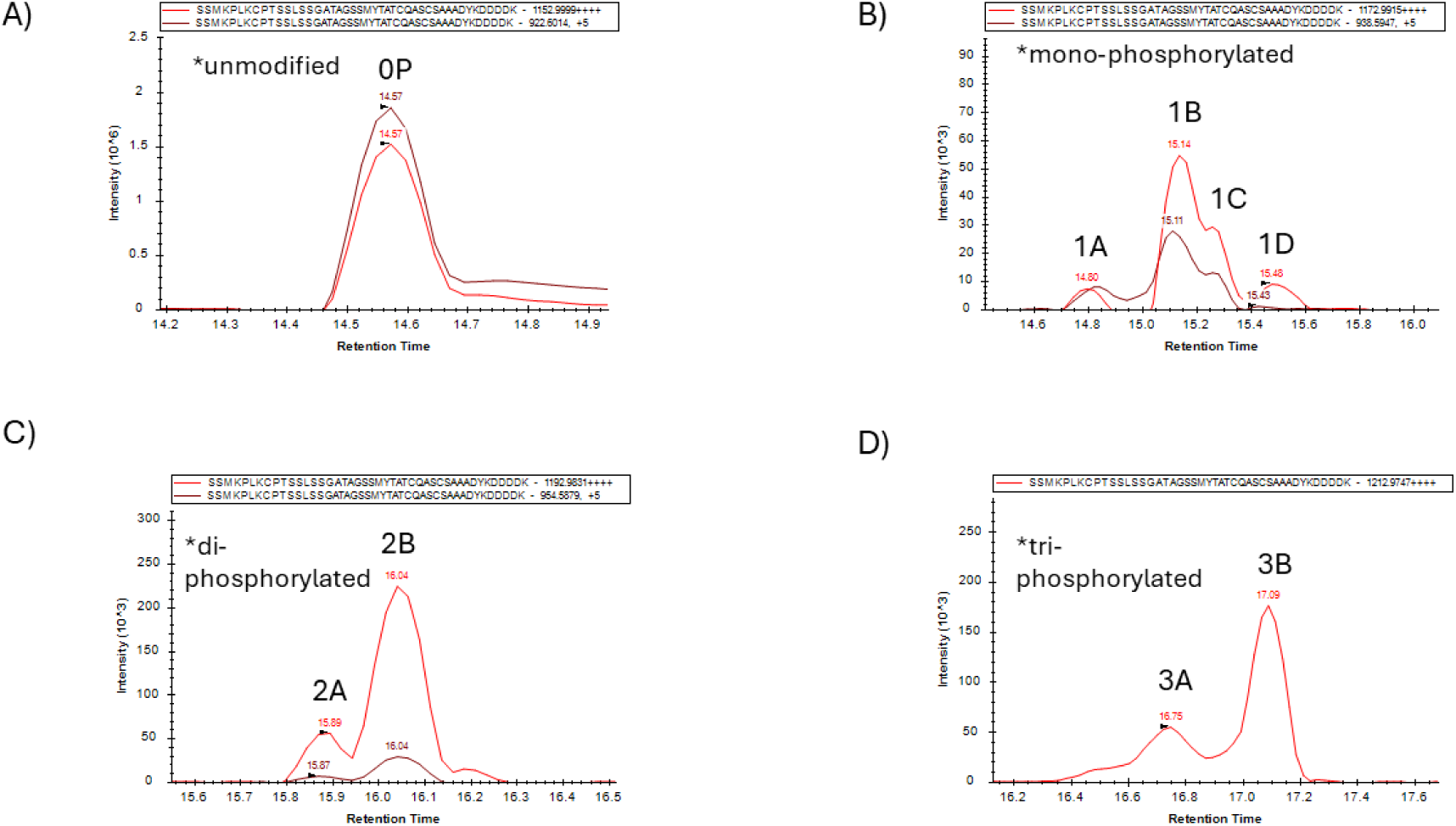
Extracted ion chromatograms of C-tail proteoforms of GLP-1R_TEV following GLP-1 treatment. The peptide analyte and m/z ratio (Th) of the precursor ions are shown in a legend above each panel. Ion intensity corresponds to the sum of the y_3_, y_4_, y_5_, y_6_, y_7_, y_8_, y_9_, y_10_ product ions. 0P= unmodified peptide; 1A-1D= four different proteoforms of mono-phosphorylated analyte; 2A-2B= two different proteoforms of di-phosphorylated analyte; 3A-3B= two different proteoforms of tri-phosphorylated analyte.

**Supplemental Figure 4.**
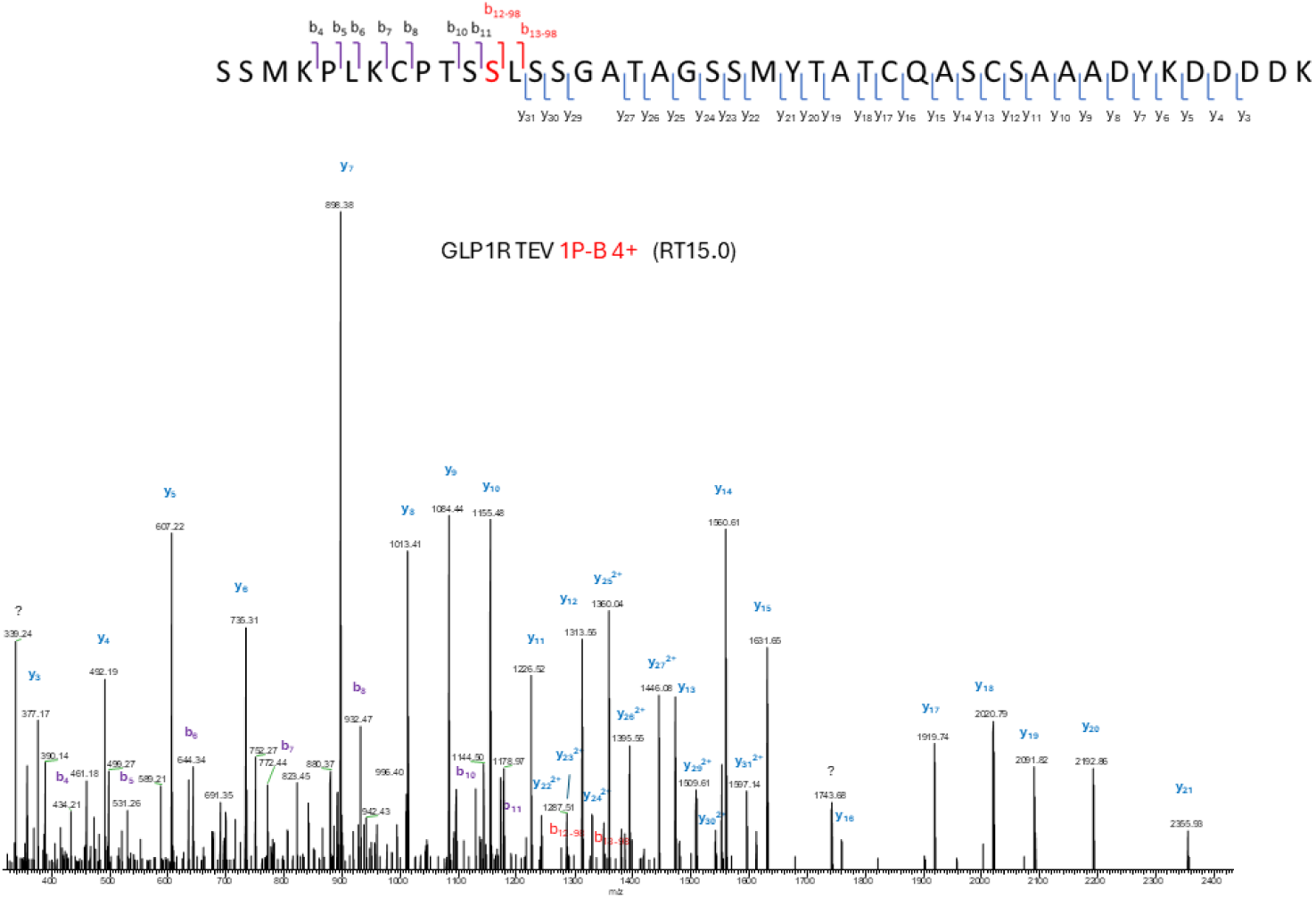
Localization of sites of phosphorylation within the GLP-1R C-tail. Sites of phosphorylation within the C-tail peptides were localized from inspection of the product ion spectra for each proteoform. For example, upon dissociation of the [M+4H]^4+^ precursor ion corresponding to GLP-1R TEV proteoform 1P-B (shown here) the MS/MS data yielded extensive b and y product ions. Location of the phosphorylation to the serine in position 12 was evidenced from the detection of (1) the non-phosphorylated b_11_ and y_31_ ^2+^ ions and (2) the detection of the b_12-98_ and b_13-98_ ions. For certain proteoforms we lacked adequate sequence specific product ions to localize the phosphorylation to a single amino acid. In these cases, we report the minimal possible sites of modification based on the information available.

**Supplemental Figure 5.**
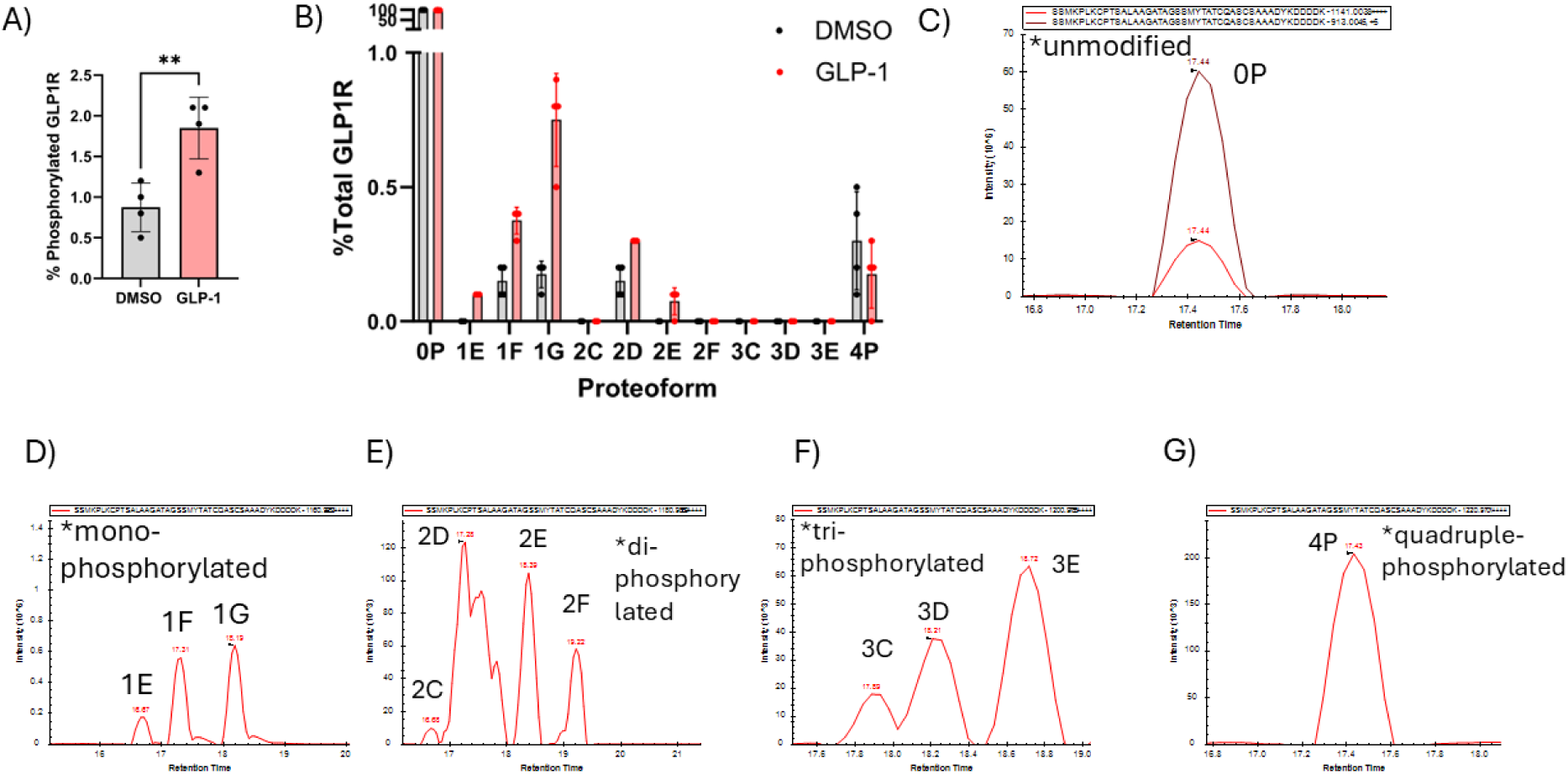
Middle-down proteomic analysis of C-tail phosphorylation in GLP-1R S442A/S444A/S445A mutant. A) Bar graph indicating the percentage of the GLP-1R S442A/S444A/S445A mutant C-tail that is phosphorylated following treatment with 1µM GLP-1(7-36). Data points and error bars are the mean and SD of technical quadruplicates from a representative example of three independent experiments. An unpaired, two-tail t-test was conducted for statistical analysis; *\*\*P* < 0.01. B) Bar graph indicating the percentage of each discrete proteoform among total GLP-1R S442A/S444A/S445A triple mutant receptor after treatment with DMSO (grey bars) or GLP-1(7-36) (red bars). 0P is an unmodified GLP-1R S442A/S444A/S445A proteoform; 1E-1G are mono-phosphorylated proteoforms; 2C-2F are di-phosphorylated proteoforms; 3C-3E are tri-phosphorylated proteoforms; 4P is a tetra-phosphorylated proteoform. C-G) Extracted ion chromatograms of the proteoforms of the GLP-1R triple mutant that are shown in supplemental Figure 4B. The peptide analyte and m/z ratio (Th) of the precursor ions are shown in a legend above each panel. Ion intensity corresponds to the sum of the y_3_, y_4_, y_5_, y_6_, y_7_, y_8_, y_9_, y_10_, y_11_ product ions.

**Supplemental Figure 6.**
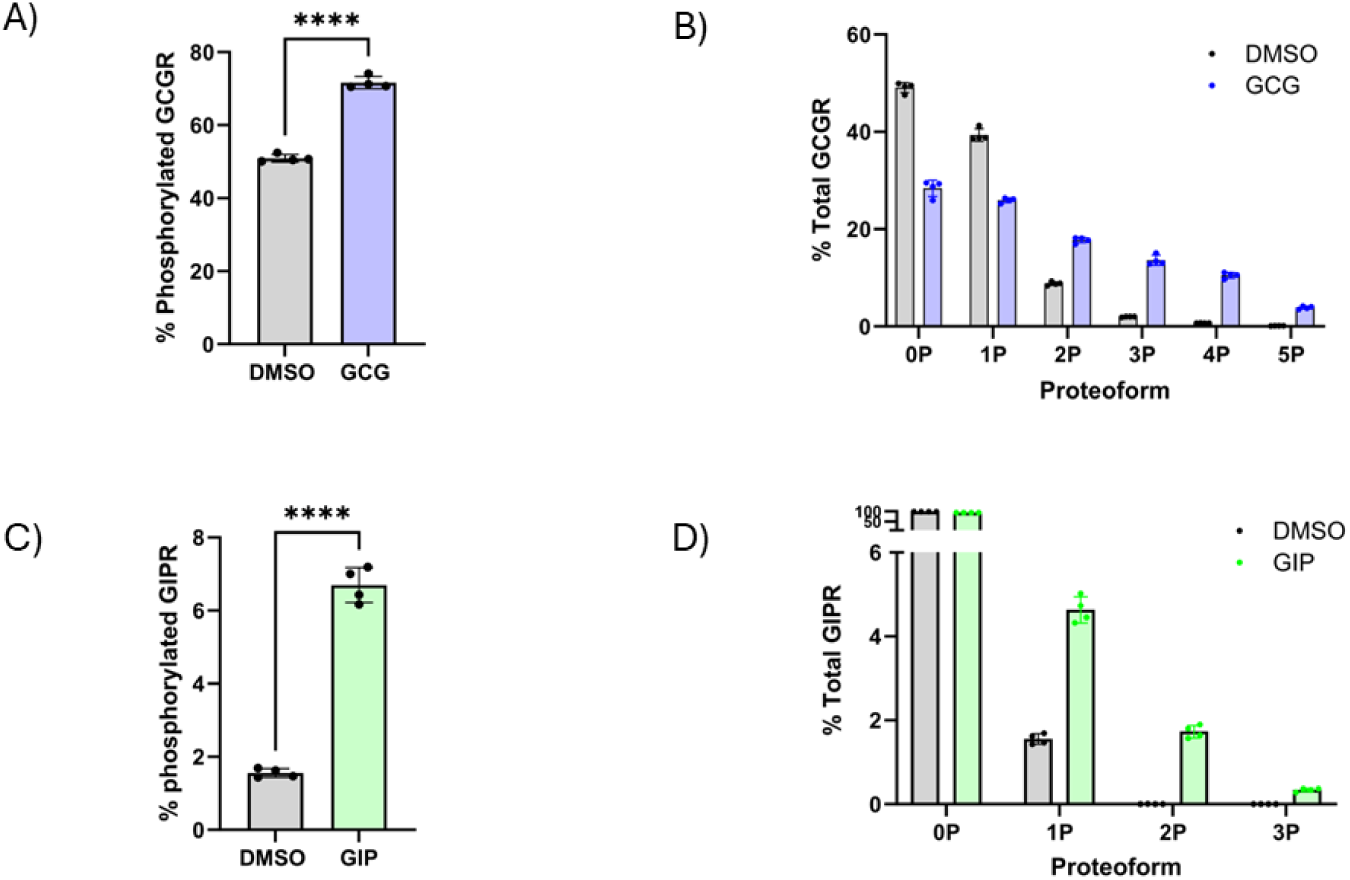
Middle-down proteomic analysis of C-tail phosphorylation in GCGR_TEV and GIPR_TEV receptors. A) Bar graph indicating the percentage of the GCGR_TEV C-tail that is phosphorylated following treatment with 1 µM GCG (1-29). B) Bar graph indicating the percentage of each discrete proteoform among total GCGR_TEV receptor after treatment either 1 µM GCG (1-29) (blue bars) or DMSO (grey bars). 0P is an unmodified proteoform; 1P is a mono-phosphorylated proteoform; 2P is a di-phosphorylated proteoform; 3P is a tri-phosphorylated proteoform; 4P is a quadruple-phosphorylated proteoform; 5P is a penta-phosphorylated proteoform. C) Bar graph indicating the percentage of GIPR_TEV C-tail that is phosphorylated following treatment with 1 µM GIP (1-42). D) Bar graph indicating the percent abundance of each discrete receptor C-tail proteoform among total GIPR_TEV receptor in cells treated with GIP (1-42) (green bars) or DMSO (grey bars). For each graph for receptor (GCGR_TEV and GIPR_TEV), data points and error bars are the mean and SD of technical quadruplicates from a representative example of three independent experiments. An unpaired, two-tail t test was conducted for statistical analysis; *\*\*\*\*P* < 0.0001.

**Supplemental Figure 7.**
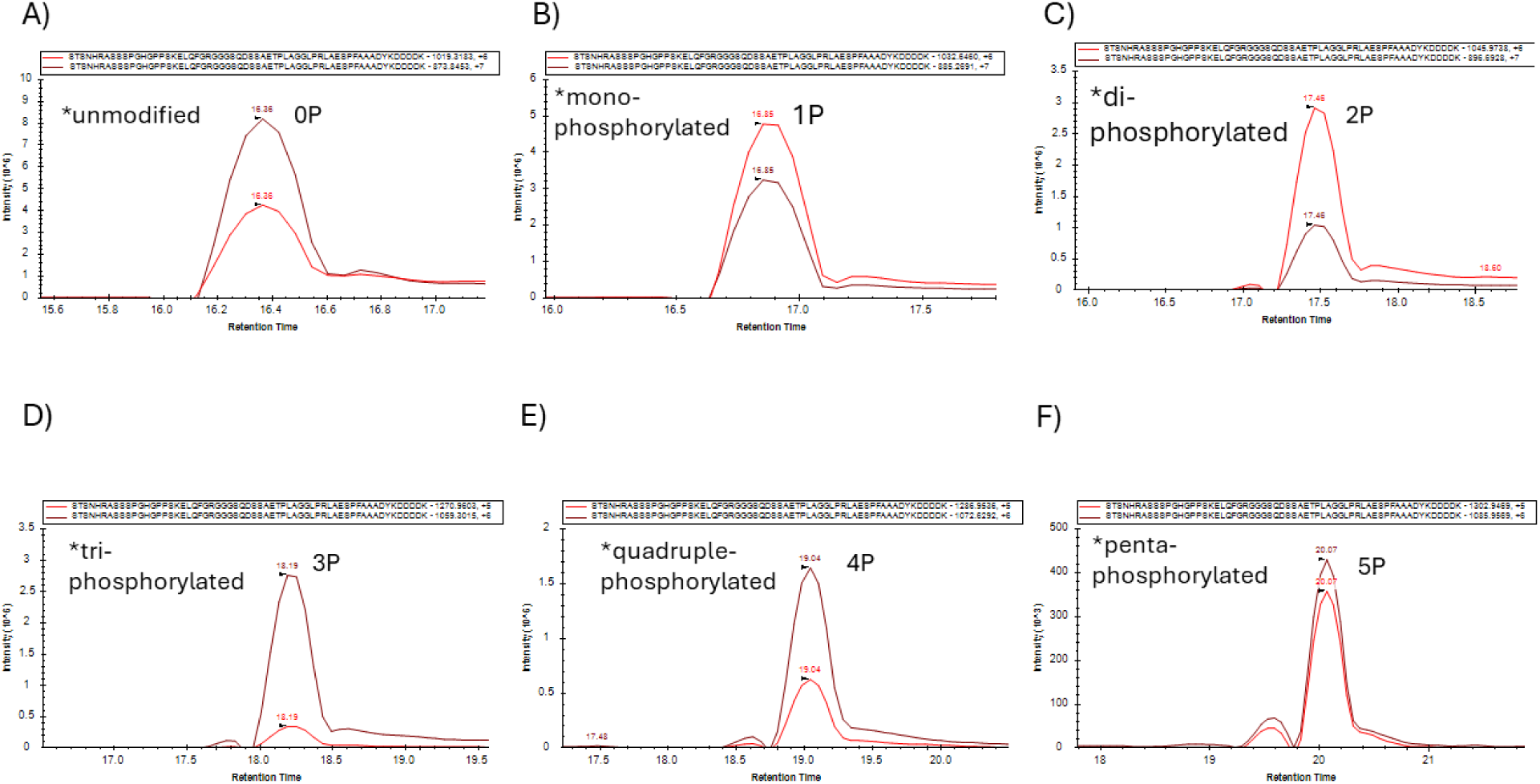
Extracted ion chromatograms of the of GCGR_TEV C-tail proteoforms following GCG treatment. The peptide analyte and m/z ratio (Th) of the precursor ions are shown in a legend above each panel. Ion intensity corresponds to the sum of the y_4_, y_5_, y_6_, y_7_, y_8_, y_9_, y_10_, y_11_ product ions.

**Supplemental Figure 8.**
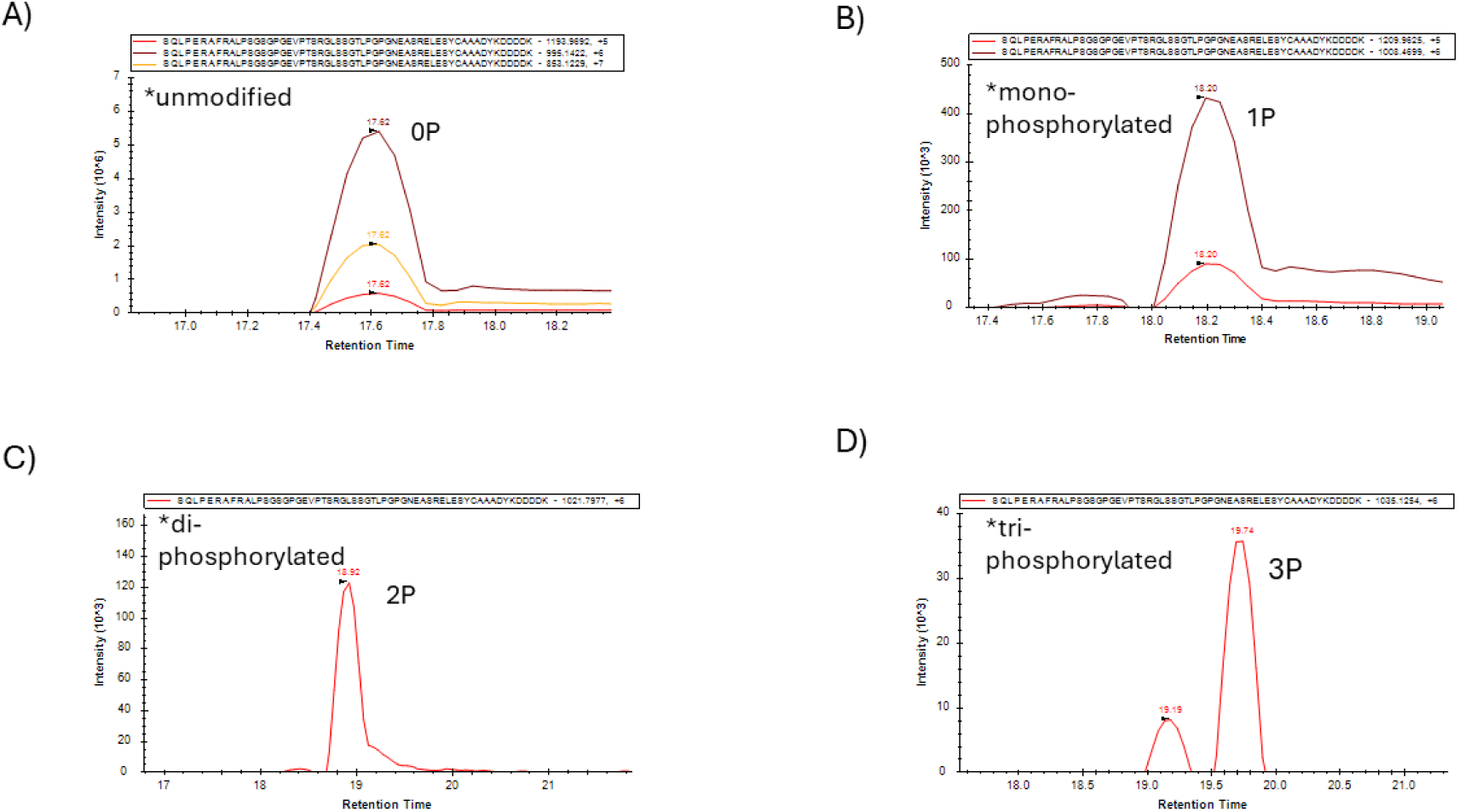
Extracted ion chromatograms of GIPR_TEV C-tail proteoforms following GIP treatment. The peptide analyte and m/z ratio (Th) of the precursor ions are shown in a legend above each panel. Ion intensity corresponds to the sum of the y_4_, y_5_, y_6_, y_7_, y_8_, y_9_, y_10_, y_11_ product ions.

**Supplemental Table 1.**
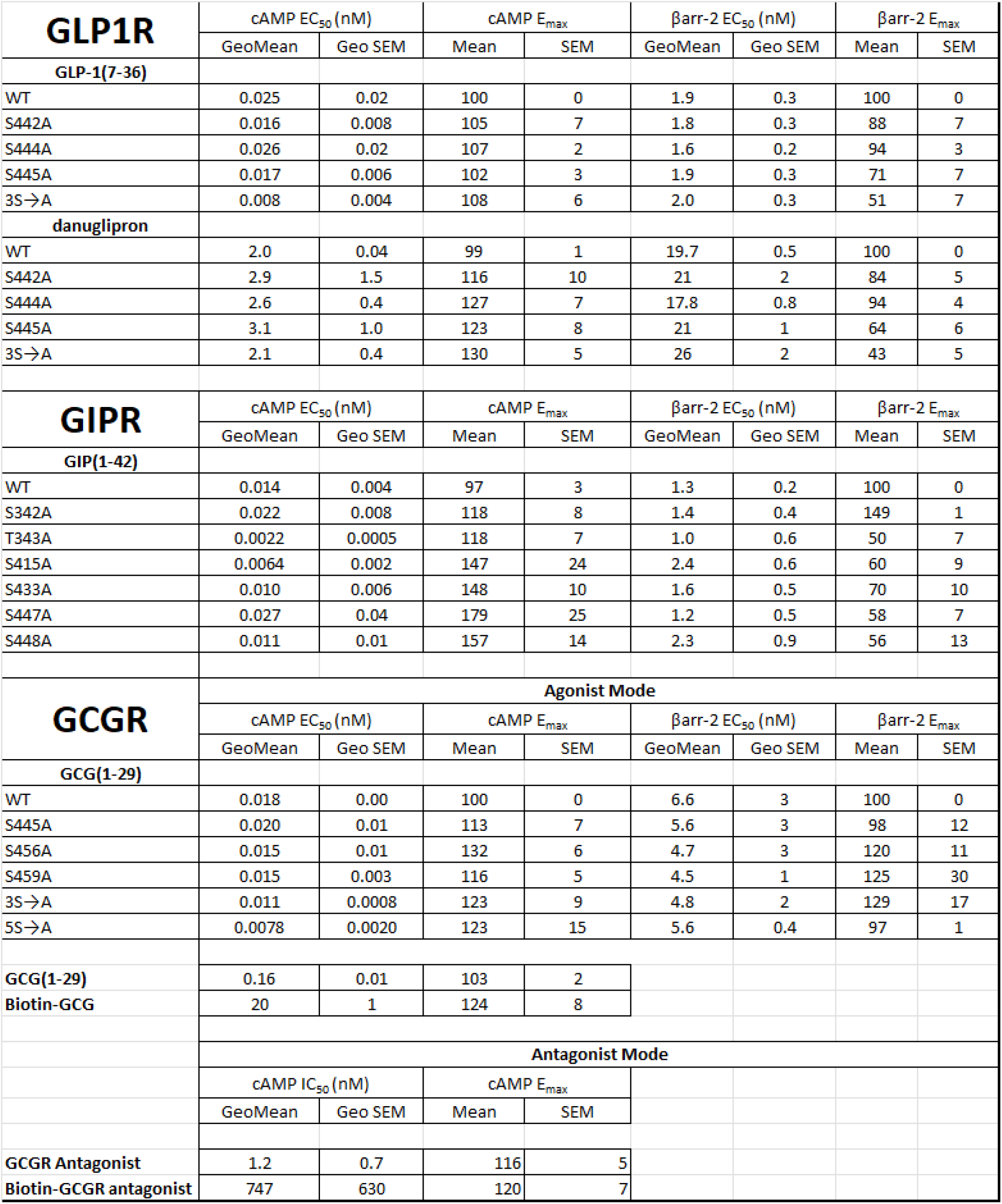
Pharmacological parameters for cAMP accumulation and β-arrestin recruitment assays. Dose response curves were fit using the 4-parameter logistic equation. Pharmacological parameters were obtained, and potency values are expressed as geometric means and standard errors and efficacy values (E_MAX_) are expressed as arithmetic means and standard errors. All data sets are generated from n=3 independent experiments.

**Supplemental Table 2.** Identities of raw mass spectrometry files uploaded to https://repository.jpostdb.org/ (jPOSTrepo).

| **<u>Experiment Number</u>** | **<u>Biological Replicate</u>** | **<u>Associated Figure</u>** |
| --- | --- | --- |
| C01089-044 | 1 | supplemental figure 1 |
| C01089-053 | 2 | supplemental figure 1 |
| C01089-054 | 3 | supplemental figure 1 |
| C01089-049 | 1 | figure 2 |
| C01089-050 | 2 | figure 2 |
| C01089-051 | 3 | figure 2 |
| C01089-042 | 1 | figure 3B and D |
| C01089-056 | 2 | figure 3B and D |
| C01089-060 | 3 | figure 3B and D |
| C01089-065 | 1 | figure 3C and E |
| C01089-066 | 2 | figure 3C and E |
| C01089-022 | 1 | figure 1 |
| C01089-031 | 2 | figure 1 |
| C01089-036 | 3 | figure 1 |
| C01089-008 | 1 | supplemental figure 2 |
| C01089-055 | 1 | supplemental figure 5 |
| C01089-065 | 2 | supplemental figure 5 |
| C01089-065 | 3 | supplemental figure 5 |
| C01089-059 | 1 | supplemental figure 6A and B |
| C01089-059 | 2 | supplemental figure 6A and B |
| C01089-061 | 3 | supplemental figure 6A and B |
| C01089-058 | 1 | supplemental figure 6C and D |
| C01089-058 | 2 | supplemental figure 6C and D |
| C01089-062 | 3 | supplemental figure 6C and D |

